# scShapes: A statistical framework for identifying distribution shapes in single-cell RNA-sequencing data

**DOI:** 10.1101/2022.02.13.480299

**Authors:** Malindrie Dharmaratne, Ameya S Kulkarni, Atefeh Taherian Fard, Jessica C Mar

## Abstract

**Background:** Single cell RNA sequencing (scRNA-seq) methods have been advantageous for quantifying cell-to-cell variation by profiling the transcriptomes of individual cells. For scRNA-seq data, variability in gene expression reflects the degree of variation in gene expression from one cell to another. Analyses that focus on cell-cell variability therefore are useful for going beyond changes based on average expression and instead, identifying genes with homogenous expression versus those that vary widely from cell to cell.

**Results:** We present a novel statistical framework *scShapes* for identifying differential distributions in single-cell RNA-sequencing data using generalized linear models. Most approaches for differential gene expression detect shifts in the mean value. However, as single cell data are driven by over-dispersion and dropouts, moving beyond means and using distributions that can handle excess zeros is critical. *scShapes* quantifies gene-specific cell-to-cell variability by testing for differences in the expression distribution while flexibly adjusting for covariates if required. We demonstrate that *scShapes* identifies subtle variations that are independent of altered mean expression and detects biologically-relevant genes that were not discovered through standard approaches.

**Conclusions:** This analysis also draws attention to genes that switch distribution shapes from a unimodal distribution to a zero-inflated distribution and raises open questions about the plausible biological mechanisms that may give rise to this, such as transcriptional bursting. Overall, the results from *scShapes* helps to expand our understanding of the role that gene expression plays in the transcriptional regulation of a specific perturbation or cellular phenotype. Our framework *scShapes* is incorporated into Bioconductor R package (https://github.com/Malindrie/scShapes).

## Background

Variation in gene expression from one cell to another plays an instrumental role in how tissues develop and function [1], and therefore consideration of the distribution shape of a gene’s expression profile is important for understanding transcriptional regulation [2]. The analysis of scRNA-seq data sets has provided a means to quantitatively study cell-to-cell variation in response to phenotypic changes and as a result, new rare and complex cell populations have been identified [3], regulatory relationships among genes have been discovered [4], and trajectories of cell lineages in development and disease have been elucidated [5]. However, a major limitation of existing scRNA-seq methods is that these analyses tend to focus on detecting changes in average expression rather than changes in variability or other properties of the distribution. While transcriptional regulation requires both, a significant strength of scRNA-seq data is the opportunity to model gene expression variability and its role in regulating cellular phenotypes.

Existing scRNA-seq methods assume that a single parametric distribution is adequate for modelling the shape of a gene’s expression profile. While this assumption may have practical advantages, it does limit the ability to discover new relationships or gene expression shape patterns. A property of scRNA-seq data is its increased sparsity compared to bulk gene expression data, where an abundance of cells may have unobserved expression levels due to either biological or technical sources of cell-to-cell variability [6, 7]. As a result, it may be an over-simplification to assume that the expression profiles of genes across the transcriptome may be adequately explained by a single parametric distribution. A limitation of this assumption may be that some genes are overlooked or modelled sub-optimally as we fail to first evaluate the prevalence of different distributions in a scRNA-seq dataset [2].

To model the heterogeneity of gene expression data under a statistical framework, it is vital that the distribution with the most appropriate fit for each gene’s expression profile be used [8]. While some statistical methods have appealed to the use of mixture models [6, 9] as an alternative to the widely-used negative binomial distribution [10, 11], they fail to investigate the range of different gene expression distributions that may be present in the scRNA-seq data as a first step. In *scShapes*, we address this issue directly by proposing a gene-specific framework for identifying differential distributions from a selection of relevant candidate distributions.

Currently differentially distributed genes can be assessed using the *R* package *scDD* [12]. However, this approach is unable to adjust for covariates directly in the model. Because accounting for covariates, especially technical ones like gender or hospital site, may be necessary to address for the underlying question being asked of the scRNA-seq analysis, this limitation of *scDD* has practical implications as the size and complexity of scRNA-seq datasets grow. The *scDD* method also only permits pairwise comparisons between two biological conditions. In our *scShapes* framework, we propose an approach for modelling read counts from droplet-based scRNA-seq experiments using generalized linear models (GLM) with error distributions belonging to the family of zero-inflated negative binomial distributions.

Zero counts are a prevalent feature of scRNA-seq data that present challenges for statistical modelling, mainly because these zero values result from different zero generating processes within the same biological system [13]. Because these zeroes can arise from biological and technical sources, the treatment of the zero values, e.g. through imputation or filtering, is not straightforward. The zero values observed in scRNA-seq can be the result of limitations in sequencing depth and capture efficiency, such that only a small percentage of the transcripts present end up being counted and low abundance transcripts can go undetected [14, 15]. Observing a zero count can also be due to biological factors like a gene that is simply not expressed in a specific cell type or state or the fact that transcription is a stochastic process which gives rise to these additional zeros [16]. By taking all these factors into account, we assume that the overabundance of zeros in scRNA-seq data is gene-specific and that each gene therefore requires a statistical model with its own set of distributional assumptions. To implement this, our framework models gene expression data for a mix of unimodal and zero-inflated distributions to identify the distribution that provides the most appropriate fit for each gene (Figure 1).

**Figure 1:**
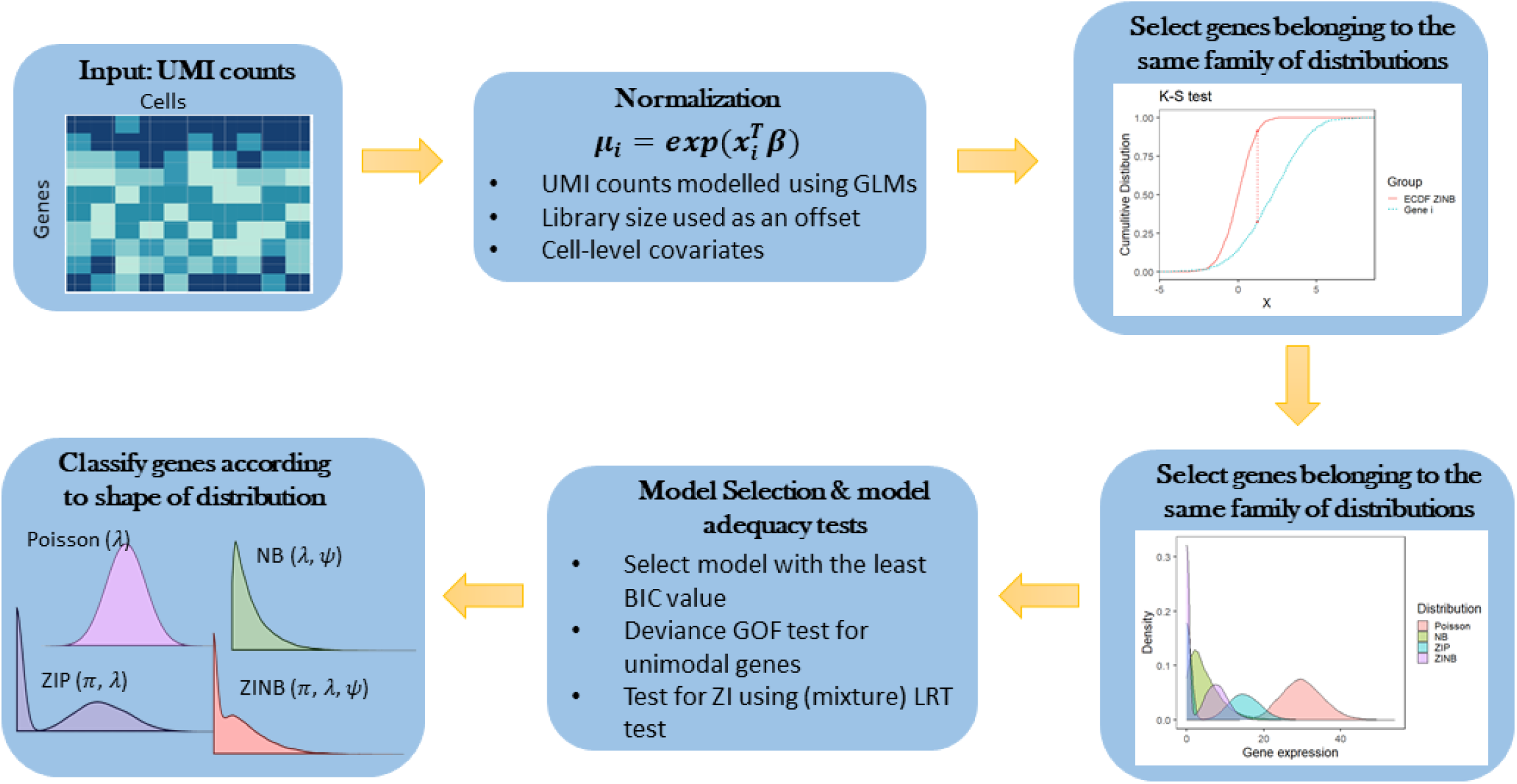
ScShapes pipeline for identifying gene expression according to distribution shape. The input to *scShapes* is a matrix of UMI counts with initial quality control performed to remove lowly-expressed genes. A Kolmogorov-Smirnov test is first performed to select genes belonging to the family of ZINB distributions. Genes that do not belong to this category are removed from further analysis. For genes that are assumed to follow a distribution from the ZINB family, the best distribution that explains most of the heterogeneity in its gene expression values is selected from Poisson, NB, ZIP, ZINB distributions based on two rounds of model adequacy testing.

Here we develop a framework, *scShapes*, which relies upon modelling read counts using GLMs and matches a distribution shape to a gene’s expression distribution for an individual gene. The gene expression distribution is modelled using both unimodal and zero-inflated distributions, where for accurate evaluation of zero-inflation we conduct a modified likelihood ratio test (LRT) [17]. Our framework has been designed to identify subtle variations from one cell to another that is not weighted towards a change in mean. It also has the flexibility to adjust for covariates and perform multiple comparisons between sample or treatment groups while explicitly modelling the variability of gene expression occurring between cells. Using simulation studies, we show that *scShapes* can reliably detect zero-inflated genes, and when applied to a range of scRNA-seq published datasets, *scShapes* can identify genes and pathways linked to the phenotype of interest that were not discovered through standard analyses of transcriptomic data. The flexibility of *scShapes* is also demonstrated through use cases that have been applied to scRNA-seq datasets that feature distinct experimental designs and experimental models.

## Data Description

A collection of 3 publicly available scRNA-seq datasets were used in this study. These datasets were downloaded from NCBI GEO (accession numbers and data descriptions given under ‘Datasets used’). For all the datasets cell-type identification has been performed by their respective original publications, which we have used in the *scShapes* framework as prior biological knowledge. For each dataset, genes with non-zero expression in at least 10% of all cells within a treatment condition are retained and were used for all subsequent analyses.

## Analyses

### An overview of the *scShapes* framework

The entry point to the *scShapes* pipeline is a set of aligned read counts from a scRNA-seq experiment. For each treatment condition, we model a gene independently using the error distributions from all four possible distributions, Poisson, Negative Binomial (NB), Zero-inflated Poisson (ZIP) and Zero-inflated Negative Binomial (ZINB) with a log link function. A model-based normalization is applied to the read counts to account for differences in sequencing depth between libraries. This is done by including the log10 of the total UMI counts assigned per cell as an offset in the GLM model so that differences in sequencing depth are directly adjusted for. Furthermore, additional covariates can be incorporated in the GLM framework to account for any biases introduced by biological replicates or technical attributes of the experimental design like batch effects. The most appropriate model is first selected based on the Bayesian Information Criterion (BIC) and LRT statistic, to ensure the best distribution is selected for each gene (see Figure 1 and Methods for more details).

### The overabundance of zeros in scRNA-seq data is gene-specific

To demonstrate the *scShapes* differential distribution framework, we apply this to a scRNA-seq dataset collected for an ageing study on two tissues, adipose and muscle, from mice (Figure 2a). In the study design, there are three groups of mice which we designate as Young, Old and Treated, where each group consists of four male mice as biological replicates (see Methods). GLM models with error distributions from the Poisson, NB, ZIP and ZINB distributions are fitted without assuming any prior biological knowledge and correcting only for the technical variability between cells. This was done by including an offset in the GLM model to account for differences in sequencing depth and the mouse ID included as an explanatory covariate in the GLM to account for any biases introduced due to one mouse being potentially different from any of its other biological replicates.

**Figure 2:**
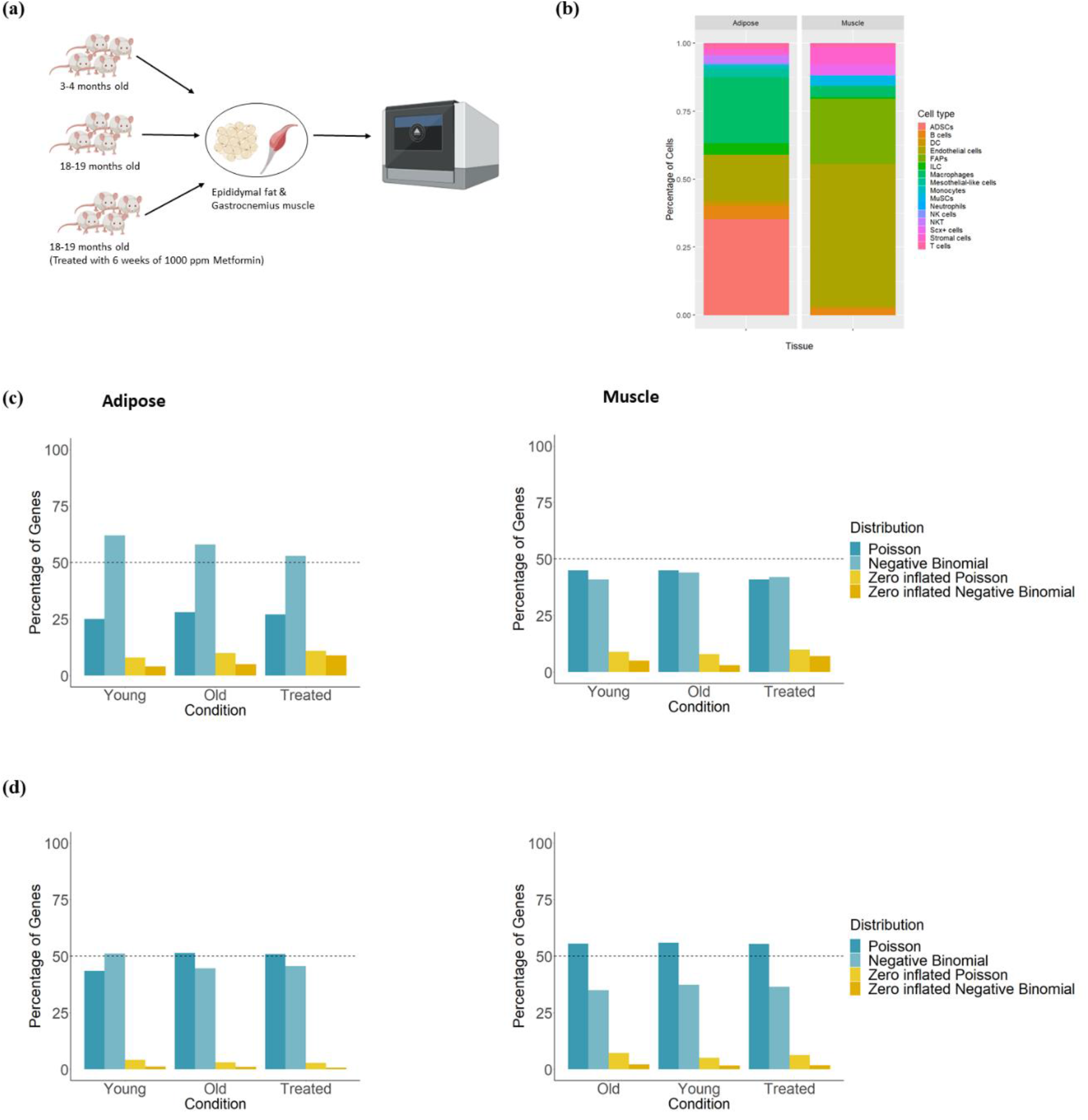
**(a)** Overview of the study design for the single-cell RNA-sequencing dataset using adipose and skeletal muscle in aging and metformin-treatment in mice. Single cells have been isolated from epididymal fat and gastrocnemius muscle of four male C57BL/6J mice in each of the three groups old (18-19 months), young (3-4 months) and metformin-treated for 6-weeks (18-19 months). Single cell library preparation was done using the 10X Chromium single cell 3’ v2 kit, with quality control and gene expression quantification carried out in Cellranger. **(b)** Cell type abundances of the single-cell RNA-seq data in adipose and skeletal muscle. Seurat R package has been used for clustering and differential expression. Cells have been annotated manually by inspecting the gene expression profiles of marker genes identified using *FindAllMarkers* in Seurat for each cluster and validated using a self-built Garnett cell type classifiers. The cell type membership was used as an explanatory covariate in the GLM framework of *scShapes*, to determine whether using prior biological knowledge would change the composition of genes following one of the four distributions.

We found that the majority of the genes (between 52%-62% in adipose and between 40%-45% in muscle) follow a NB distribution in both tissue types (Figure 2c). In the muscle tissue, it is worth highlighting that at least 50% of the genes do not follow the NB distribution under the three treatment conditions. Collectively, over 55% of genes follow either a NB, ZIP or ZINB distribution and this observation aligns with the nature of scRNA-seq data which is driven by over-dispersion and zero-inflation. These results provide evidence that in fact not all genes have expression profiles that follow a NB distribution and more importantly, demonstrate that not all the genes in the transcriptome follow a single distribution either.

### The inclusion of prior biological knowledge, like cell type membership, changes the shape of a gene’s expression distribution

We investigated whether accounting for known biological knowledge such as the membership of a specific cell type affects the classification of genes into their distribution shape. The ageing mouse dataset consists of 13 known cell types in adipose and 12 known cell types in muscle (Figure 2b). To account for the biological variation in the data, we introduced the cell type information as an explanatory covariate in the GLM model. Accounting for cell type membership resulted in a marked reduction in the number of NB-distributed genes, a resulting increase in Poisson-distributed genes, and the number of zero-inflated genes also decreased (Figure 2d & Table S1). This observation is in line with Choi *et al*. [18], which also observed that adjusting for cell type resulted in a reduction of the number of zero-inflated genes.

Housekeeping genes are required for the maintenance of basic cellular functions, and hence are usually uniformly expressed in all cells with low variance. Interestingly we found that around 50% of the genes under each condition that remained Poisson-distributed even after accounting for cell type membership were previously described as having a role as a housekeeping gene [19] (Figure 3). The overlap between the Poisson-distributed genes with and without accounting for known cell types was significantly different only in adipose (adjusted P-value < 0.01, Figure 3a). A gene which remains Poisson-distributed regardless of whether cell-type specific expression rates were taken into account are indicative of housekeeping genes as the rate of expression in these genes do not vary across cell types. Our finding indicates that a Poisson error model is sufficient to explain the variability in the expression of genes with a housekeeping function.

**Figure 3:**
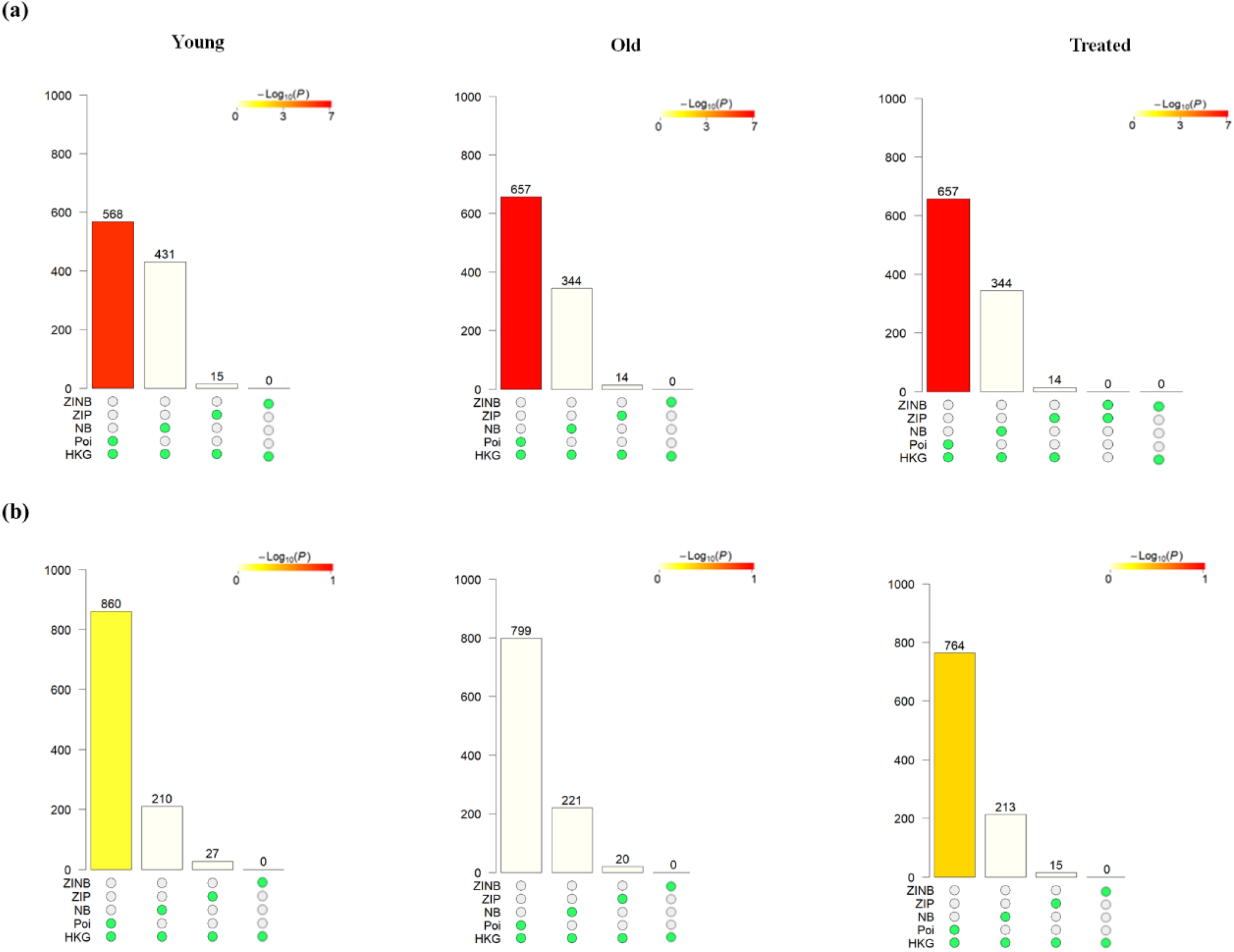
Investigating the prevalence of housekeeping genes (HKG) amongst genes that remain P, NB, ZIP, and ZINB both with models correcting for technical variability and when accounting for known biological variability in (a) adipose and (b) muscle. *scShapes* modelling framework is first applied to identify the distribution shape in old, young and metformin treated group only accounting for the technical variability in the GLM model (i.e. only including the offset term and mouse ID as a covariate in the GLM to account for differences in sequencing depth and biological replicates respectively). The same process is repeated again including information on the cell-types as a covariate in the GLM (in addition to the offset and mouse ID) to account for known sources of biological variability. We next checked for the overlap of genes that followed the same distribution in both instances above and checked for the statistical significance of the overlaps using the R package *SuperExactTest*. The height of each bar corresponds to the number of overlapping genes and the colour corresponds to the log_10_(P-adjusted) value that assess the significance of the overlap.

### A simulation study to evaluate the sensitivity of the *scShapes* gene expression distribution classification

We designed a simulation study to evaluate our framework’s ability to accurately classify genes into Poisson, NB, ZIP and ZINB distributions. To derive a realistic set of model parameters, we first performed model classification using the *scShapes* framework for the well-known 3k-cell PBMC dataset. The peripheral blood mononuclear cells (PBMCs) were downloaded from the 10X Genomics website (https://www.10xgenomics.com/resources/datasets). Using the model classification and parameter estimation we simulated count data for three sample sizes, 2638, 3000 and 5000 cells (see Methods section). As expected, we see that as the sample size increases, the ability of our framework to correctly identify the best fit distribution increases (Figure 4). However, the more striking result is that our framework can accurately identify the correct model distribution for each gene with an accuracy of 85% or above for all four distributions across the range of sample sizes tested (Figure 4a).

**Figure 4:**
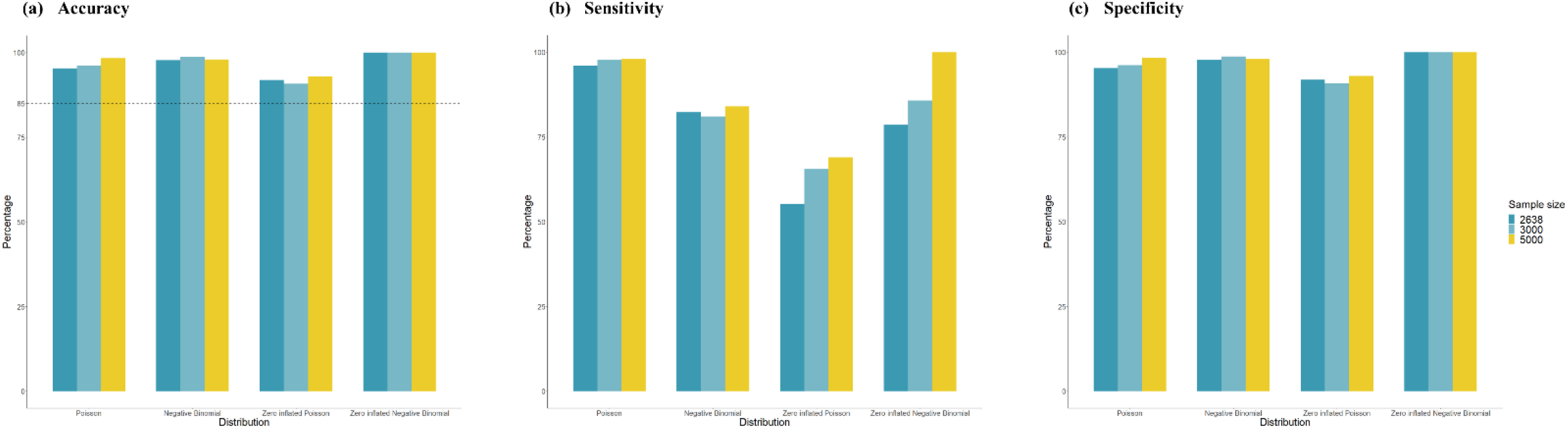
Summary statistics for the simulation study carried out to assess the performance of *scShapes* at identifying distributions of single cell gene expression data. Summary statistics (a) Accuracy, (b) Sensitivity and (c) Specificity; of the simulation study conducted to evaluate *scShapes*’s ability to classify genes into a P, NB, ZIP and ZINB distribution. Model parameters of the four distributions were estimated using the 3k PBMC dataset and were used for simulating gene expression values from P, NB, ZIP, ZINB distributions using three sample sizes *n* (i.e. number of single cells). Next, *scShapes* was run on the simulated gene expression counts to identify the distribution of the simulated data. Results of the simulation study was evaluated using the three summary statistics accuracy, sensitivity and specificity.

### The zero-inflation parameter can be interpreted as an estimate of biological zeros

Being able to distinguish the zeros that reflect technical drop-outs versus genuine biological zeros is helpful for understanding how genes control cellular phenotypes. The zero-inflation parameter 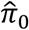 in the *scShapes* model gives an estimate of the proportion of structural zeros for each gene which may be indicative of the subset of cells where the transcript is truly absent. These models make the assumption that the zero observations have two different origins namely, **structural zeros assumed to be observed due to specific structure in the data** which are distinct from **random zeros which are assumed to be observed due to sampling variability**. In order to evaluate the relationship between the estimated zero-inflation parameter 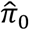 and the percentage of zeros across all cells in genes detected to be zero-inflated (*i.e*. genes following either a ZIP or ZINB distribution), we plotted the estimated zero-inflation parameter against the percentage of zeros across cells for each zero-inflated gene (Figure S1). While most features have at least 50% or more zeros across cells, the zero-inflation parameter can vary between any value between 0 and 1. This indicates that there is either no relationship or a very weak relationship between the zero-inflation parameter and the percentage of zeroes. It demonstrates that genes with a high percentage of zeroes overall are not necessarily captured by models where the zero-inflation parameter is high. Choi *et al*. [18] has shown that the primary cause of zero inflation is due to biological variation and not only due to technical variation. So, one can assume that the excessive zeros we observe in single-cell data are the result of the two zero generating processes; where either a transcript is absent from a biological system due to sampling variability or where a transcript is truly absent from the biological system. Hence, modelling genes with excess zeros using zero-inflated distributions will be ideal in such a scenario.

We also leveraged the zero-inflation parameter to identify genes that had an excessively high number of structural zeros. For example around 98% of the zeros in the gene *Arhgap20* were structural zeros in adipose under the Old condition. This gene encodes a protein which is an activator of RHO-type GTPases. It has been shown to be associated with Alzheimer’s disease and predicted to be a dysregulated gene [20]. Similarly, the gene *Sfxn1* which has around 85% of structural zeros based on the estimate of the zero-inflation parameter in muscle under the Old condition, is found to be downregulated in senescence (SeneQuest), a process that is a causal factor for aging tissues [21]. *Sfxn1* is a protein-coding gene which mediates the transportation of serine to the mitochondria. Although the mitochondrial biology of sideroflexin (SFXN) family are not entirely explored, dysfunction in mitochondrial carriers results in perturbations in oxidative phosphorylation which underlies various pathologies including aging related diseases [22]. Hence it could be hypothesized that the higher number of structural zeros in *Sfxn1* could be as a result of aging associated mitochondrial dysfunction with aging in the muscle. These results highlight the possibility that the estimates of the zero-inflation parameter might be indicative of features which are truly absent from a biological system and therefore play a role in the regulation of the cellular phenotype under study.

### Around 30% of the genes undergo change in shape of distribution with aging or treatment, and collectively these differentially-distributed genes demonstrate a significant enrichment of aging-related pathways

Since we observed a considerable change in the composition of unimodal genes to zero-inflated genes after accounting for known cell types, this suggests that accounting for cell type-specific effects is important for addressing biological information about distributional shapes. To investigate the biological inference from enriched pathways of differentially distributed genes, we performed pathway over-representation analysis on the model distribution classifications obtained after taking into account cell type-specific gene expression for the mouse dataset by adjusting for cell type as a covariate. We identified the genes that switched their distributions either between Old vs Young or Old vs Treated. Nearly, 30% of the genes were differentially distributed in at least one of the comparisons (Old vs Young or Old vs Treated) in both tissues adipose and muscle. The top 10 Hallmark pathways and KEGG pathways most over-represented among the genes switching distributions are represented in Figures 5 & S2 respectively.

**Figure 5:**
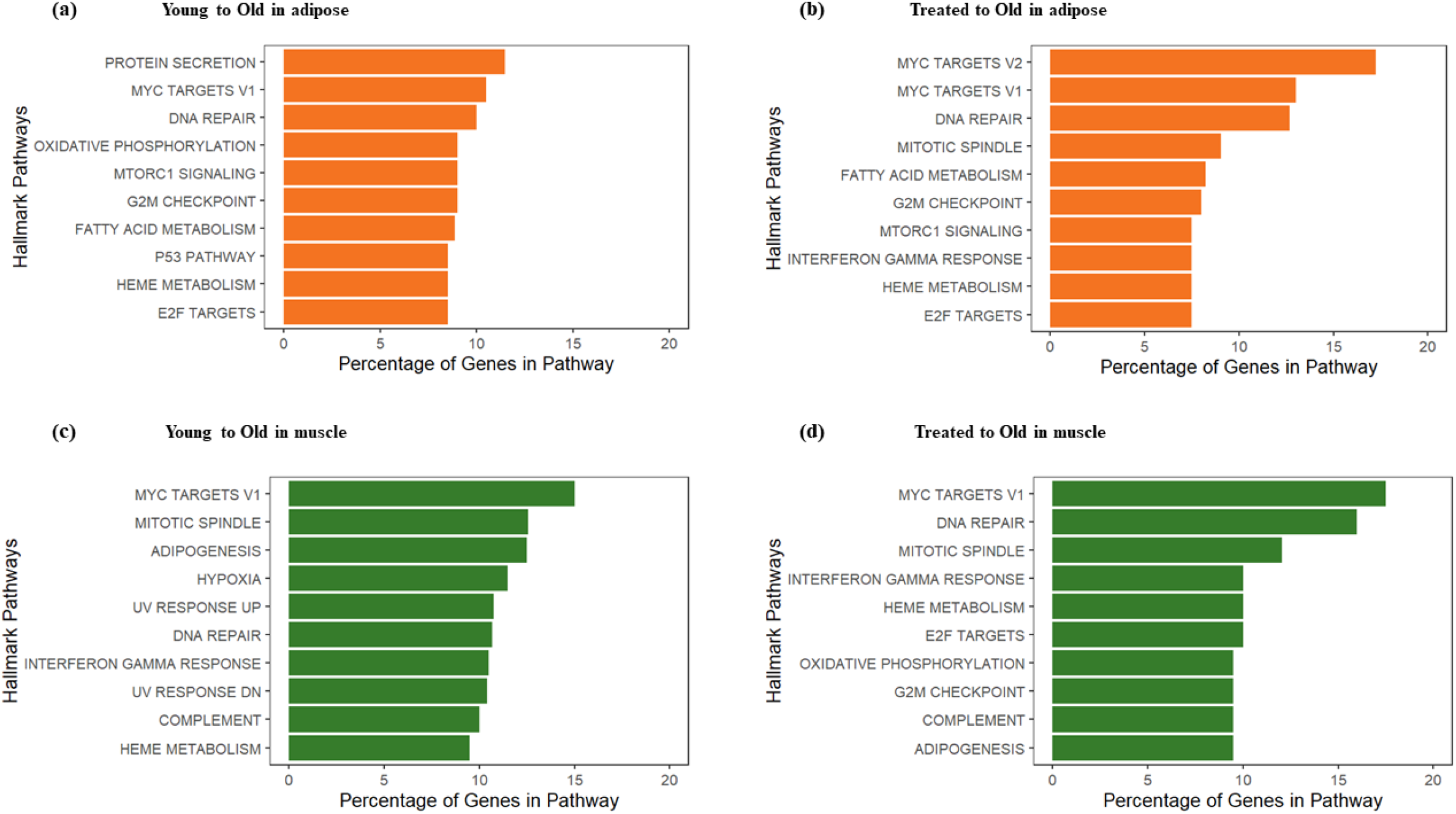
Pathway over-representation analysis of differentially-distributed genes detected by scShapes. The top 10 significant pathways (BH adjusted P-value < 1 × 10^−4^) in the pairwise comparisons (a) Young to Old in adipose (b) Treated to Old in adipose (c) Young to Old in muscle (d) Treated to Old in muscle using Hallmark pathways. The length of the bar corresponds to the percentage of differentially distributed genes that were represented in the Hallmark gene set.

The detection of statistically significant pathways provides evidence that the differentially distributed genes detected by our *scShapes* framework are enriched for biological processes related to aging. Consideration of what these processes are help to illustrate how these genes may be participating in processes that are altered through aging. For example, in the Hallmark pathways, one of the most commonly enriched most significant pathways include *DNA damage*, where excessive DNA damage or poor DNA repair contributes to the aging process [23]. Among the most enriched KEGG pathways in the Treated group in adipose is the *insulin signalling pathway* which is one of the primary nutrient-sensing pathways targeted by metformin [24]. Furthermore, the *MAPK signalling pathway*, is involved in the regulation of differentiation, cell-growth, proliferation and apoptosis [25] and is enriched in both adipose and muscle in the treated cells. Metformin is found to be relevant in activating the MAPK signalling pathway and increases the expression of DNA damage and growth inhibition gene *GADD153* [26]. Among the differentially distributed genes we find the transcription factors (TFs) *FOXO3* involved in regulating aging-associated stress response and proposed to be mediated by metformin [27, 28], *RXRA*, overexpression of which reduces DNA damage accumulation leading to delays in replicative senescence [29] and previously shown to be targeted by metformin in human adipose [30], to be common in both Old vs Young and Old vs Treated comparisons in adipose. Similarly some of the TFs commonly differentially distributed in muscle in both Old vs Young and Old vs Treated include *SRF* reduction of which leads to premature aging in skeletal muscle [31] and *IRF3* a novel inhibitor of cellular senescence and inducer of cell growth inhibition [32].

### Genes switching to zero-inflated distributions in the old condition may point to those genes involved in the regulation of transcriptional bursting

According to Clivio *et al*. [33] zero-inflated genes may reflect other biological phenomenon such as transcriptional bursting. We investigated this hypothesis in the context of aging using the metformin-treated data set. With age, a loss of transcriptional regulation is accompanied by an increase in transcriptional noise [34]. As transcriptional bursting leads to increased levels of gene expression noise, we used *scShapes* to identify the set of genes that switched from a unimodal distribution in the Young group to a more heterogeneous pattern captured by the zero-inflated distribution in the Old group while also switching from a unimodal distribution to a zero-inflated one in the Treated to Old groups respectively (see Supplementary Figure S3). The rationale behind using the *scShapes* framework to detect this specific profile is that these genes become more heterogeneous in the presence of ageing (Young to Old) as well as in response to metformin-treatment (Treated to Old). Interestingly the top Hallmark enriched term for genes switching to zero-inflated distributions in old from both young and treated in muscle is *WNT beta catenin signalling* (Figure S3), a signaling pathway that may play a role in modulating gene expression noise [35]. Some of these genes include *HDAC5*, Histone Deacetylase which plays an important role in transcriptional regulation and is involved in the chromatin dynamics of generating gene-specific noise [36]. These results draw our attention to the possibility that zero-inflated genes might be indicative of basic biological phenomena like transcriptional bursting. The use of the *scShapes* framework represents a methodological approach to detect this kind of unimodal to zero-inflated differential switching that may be useful for identifying further genes involved in the transcriptional bursting process. It is worth emphasizing that the flexibility of the *scShapes* framework and its range of distribution assumptions are what led to the identification of these genes, which would not have been necessarily detected by methods where a NB distribution is applied or assume the prevalence of a single distribution shape for all genes.

### Comparison of *scShapes* with existing tools for differential gene expression analysis

To compare the performance of our *scShapes* framework for discovering age-related gene expression to that of standard differential expression (DE), we carried out standard DE using *edgeR* [37] and also compared our framework against *scDD* [12] which models the change in mean expression levels by comparing gene expression distributions. *edgeR* was used to test for differential expression and the pairwise comparisons tested were between the Young vs Old and Old vs Treated groups. We applied *edgeR*’s quasi-likelihood approach (QLF) with cellular detection rate as a covariate, since this method was shown to perform well for DE analysis of scRNAseq data after filtering out lowly-expressed genes [38]. Genes were corrected using the Benjamini & Hochberg (BH) correction and only genes that had a corrected *P-value* of < 0.01 were retained. Although there were similar number of DE genes between *scShapes* versus *edgeR* and *scDD*, there is little overlap even between *edgeR* and *scDD* methods (Table 1). The genes found by our *scShapes* method that are exclusively switching between a unimodal distribution and a zero-inflated distribution represent over 25% of the genes in adipose and over 45% of the genes in muscle. Some of the unique genes detected by our pipeline include the TFs *FOXO3, PKNOX1* involved in apoptosis, regulation of oxidative stress and DNA damage [39] in adipose. Similarly, the TFs *IRF3, SRF, TFE3* involved in autophagy are some of the uniquely detected genes by our pipeline in muscle.

**Table 1:** Comparison of genes identified by scShapes with two other methods, edgeR and scDD. The total number of genes for *scDD* includes genes detected using the categories DE, DP, DB, and DZ.

| Tissue | Comparison | Number of Genes Detected by *scShapes* | Number of Genes Detected by *edgeR* | Number of Genes Detected by *scDD* | Overlap Between All 3 Gene Lists | Overlap between *edgeR* and *scDD* | Number of Genes Unique to *scShapes* |
| --- | --- | --- | --- | --- | --- | --- | --- |
| Adipose | Old vs Young | 902 | 2045 | 2291 | 102 | 796 | 402 |
|  | Old vs Treated | 825 | 916 | 1214 | 17 | 183 | 605 |
| Muscle | Old vs Young | 988 | 519 | 338 | 9 | 35 | 840 |
|  | Old vs Treated | 986 | 393 | 319 | 0 | 24 | 877 |

Of the genes that were detected as differentially distributed by our *scShapes* method, between 25% to 27% of the genes switch distributions from a unimodal to a zero-inflated distribution in adipose (Table S2a). However, in muscle around 50% of the differentially distributed genes switch distribution from a unimodal to a zero-inflated distribution. The genes switching distribution from a unimodal to a zero-inflated distribution are indicative of the genes that have excessive zeros in one condition compared to another condition. We further investigated the patterns of differential distributions to identify what percentages of genes switched distribution from NB to either Poisson, ZIP or ZINB (or vices versa) and this information is summarized in Table S2b. It can be seen that around 50% to 75% of the genes switched distribution between a NB to Poisson distribution (or vice versa) in both pairwise comparisons for both tissues.

We next looked at what proportion of these genes were also detected as differentially expressed by *edgeR*. Although there was not a high overlap between the differentially distributed genes detected by our pipeline and the differentially expressed genes detected by *edgeR*, we observed that most of the genes commonly detected by these two methods switch distribution from NB to Poisson, ZIP or ZINB (or vice versa, Table S3). Further investigation of the overlap of genes detected by *edgeR* and our *scShape* framework revealed that most of these genes follow a unimodal distribution in the two pair-wise comparisons in both adipose and muscle (48% to 77% of the genes, see Table 2). This is to be expected since *edgeR* detects differential expression by assuming that the gene expression levels follow a negative binomial distribution for all genes.

**Table 2:** Break down of distribution shapes of genes that were detected to be both differentially distributed by scShapes and differentially expressed by edgeR.

| Tissue | Distribution | Old vs Treated |  | Old vs Young |  |
| --- | --- | --- | --- | --- | --- |
|  |  | Old | Treated | Old | Young |
| Adipose | Poisson | 55 | 59 | 155 | 110 |
|  | NB | 56 | 51 | 113 | 137 |
|  | ZIP | 14 | 13 | 29 | 44 |
|  | ZINB | 0 | 2 | 4 | 10 |
| Muscle | Poisson | 29 | 42 | 35 | 41 |
|  | NB | 36 | 24 | 39 | 37 |
|  | ZIP | 20 | 19 | 29 | 23 |
|  | ZINB | 4 | 4 | 3 | 5 |

Similarly, we also compared the distribution patterns of commonly differentially distributed genes that were detected by our DD pipeline and that of *scDD*. Compared to *edgeR*, there is little overlap between the genes detected by our pipeline and that of *scDD* in muscle, although this number is quite similar for adipose (33% and 15% in Old vs Young and Old vs Treated respectively, see Table 3). Although both approaches are designed to detect differential distributions, they each make very different modeling assumptions about the distributions. In *scDD*, log-normalized gene expression measurements are modelled using a Bayesian non-parametric modelling framework utilizing Dirichlet process mixture models. The different assumptions that are made in *scDD* and *scShapes* mean that these methods are not equally affected by the degree of heterogeneity occurring from cell to cell in the adipose and muscle tissues, and hence different sets of genes were obtained. Of the genes that were commonly detected by *scShapes* and *scDD*, most of them were categorized either as DE or DZ (differential proportion of zeros) by *scDD*. For the genes which are not differentially distributed in the non-zero values, *scDD* checks whether the proportion of zeros are significantly different between the two conditions (DZ). We further investigated the pattern of distribution of the genes detected to be DD by our pipeline as well as classified as DZ by *scDD* (Table S4). We checked whether these genes switched from a unimodal distribution to a zero-inflated distribution between conditions. However, less than 50% of the genes would switch distribution between a unimodal distribution and a zero-inflated distribution in the genes classified as DZ by *scDD*. This suggests that *scShapes* is detecting a different type of profile change in gene expression than what can be explained by *scDD*’s DZ criteria alone.

**Table 3:** Investigating whether genes detected by scShapes share any similarities with distribution categories detected by scDD.

| Tissue | Comparison | Overlap between DD genes from *scShapes* and *scDD* | Genes detected as DB by *scDD* | Genes detected as DE by *scDD* | Genes detected as DP by *scDD* | Genes detected as DZ by *scDD* |
| --- | --- | --- | --- | --- | --- | --- |
| Adipose | Old vs Young | 301 | 3 | 177 | 0 | 121 |
|  | Old vs Treated | 112 | 2 | 47 | 0 | 63 |
| Muscle | Old vs Young | 51 | 0 | 12 | 0 | 39 |
|  | Old vs Treated | 20 | 0 | 9 | 0 | 11 |

### Additional case studies to demonstrate *scShapes*’ utility for a wide range of experimental designs

The utility of the *scShapes* framework can be best demonstrated by applying it in the context of multiple datasets that represent a diverse range of biological perturbations. The primary dataset used in this study describes one of the most common experimental designs used for scRNA-seq approaches where single cells are profiled from multiple treatment conditions, *e.g*. young, old, and treated mice. This design is illuminating for understanding ageing and metformin-induced effects on the transcriptome. It is important to recognize that other kinds of designs are represented in scRNA-seq datasets and that *scShapes* is able to infer insightful results for these too. We selected two other experimental designs to showcase the flexibility and utility of *scShapes* to identify useful biological results in the presence of different degrees of complexity present in scRNA-seq data.

### Using *scShapes* to gain insights into the transcriptional response of the immune system in COVID-19 patients

A scRNA-seq dataset was generated for peripheral blood mononuclear cells (PBMCs) from seven patients hospitalized for COVID-19 and six healthy controls [40] (Figure S4a). We selected this dataset to showcase how *scShapes* can detect differential distributions in the presence of a much higher number of cells (~44,721 cells versus ~12,987 in the previous ageing dataset) and make use of multiple covariates. In the original study, Wilk *et al*. [40] classified each cell in the COVID-19 patient data and healthy controls into one of 20 known cell types.

Cell type membership, donor ID and gender were included in *scShapes* as explanatory covariates in the GLM. *scShapes* was used to determine the prevalence of genes classified into each of the four distributions Poisson, NB, ZIP and ZINB (Table S5) and determine genes that switched distributions between the COVID-19 group and the healthy control group.

Interestingly, we found that over 80% of genes followed a NB distribution, with the number of genes Poisson distributed being negligible. Pathway over-representation analysis of the genes switching distribution between COVID-19 group and healthy controls revealed pathways like *oxidative phosphorylation*, *interferon response*, and *immune response* related pathways (Figure S4 b-d) which are all pathways previously implicated with COVID-19 [41, 42].

We also used this dataset to investigate the nature of differential distributions occurring within a specific cell type. We focused our attention on the seven main cell-types, B cells, natural killer (NK) cells, dendritic cells (DCs), CD4^+^ T cells, CD8^+^ T cells, CD14^+^ monocytes and CD16^+^ monocytes. We found that there was significant overlap between the differentially-distributed genes in NK cells and the T cells (Figure S5a). We also found that differentially-distributed genes were enriched for interferon (IFN) responses, inflammatory responses and NF-κB signaling pathway (Figure S5b). We also saw a statistically significant higher enrichment of *apoptosis*, metabolic pathway *oxidative phosphorylation* and *WNT/β-catenin signalling* between the differentially distributed genes between COVID-19 group and healthy controls in NK cells, CD4^+^ T cells and CD8^+^ T cells.

### Pathway over-representation analysis of simulated T-cells highlights pathways implicated in T-cell activation

In contrast to the previous designs, we included a design that featured a single cell type that was profiled in different tissues. This kind of experimental design is relevant to scRNA-seq datasets because we have already shown that over-dispersion and zero-inflation observed in gene expression rates are influenced by the presence of different cell-types. A study generated scRNA-sequencing data of human T-cells isolated from bone marrow (BM), lungs (LG) and lymph node (LN) from two adult deceased organ donors and blood (BL) from two healthy donors for comparison [43] (Figure S6a). Tissues acquired from the donors were stimulated resulting in over 50,000 resting and activated T-cells. We used *scShapes* to model the gene expression distributions separately under the two treatment conditions, resting (non-stimulated) and activated (stimulated) where the donor ID was used as an explanatory covariate that was adjusted for in the GLM model. As expected, the Poisson distribution was the most prevalent distribution across the four tissue types in both treatment conditions. Interestingly we see that there were more genes with distributions that were classified as either NB, ZIP or ZINB distributed across stimulated T cells in all tissue types and blood (see Table S6) compared to resting T cells. The genes that switched distributions between resting and activated (Table S7) in the four tissue types were enriched for *activation of immune response* and other GO terms (see Figure S6 b-e). This result likely reflects the fact that in BM, LG and LN, the activated T-cells secrete cytokines that regulate immune response. The differentially distributed genes in this pathway that are shared between BM, LG and LN include *PSMC1, PSMC3* where both of these genes are part of the 26S proteasome involved in cell-cycle progression, DNA damage repair, apoptosis, and *XRCC6* involved in regulation of innate immune response [44].

## Discussion

We have presented a flexible statistical framework for quantifying gene expression heterogeneity of scRNA-sequencing data by modelling gene expression levels under a differential distribution pipeline. What makes our method unique is that it is fundamentally different from a method that detects differential expression by testing for the significance of change in mean of the expression rates. Our framework identifies and classifies genes according to their shape of gene expression distribution and allows for the possibility that one gene’s expression profile across a population of single cells may be different from another gene. As scRNA-seq data are characterized by properties of sparsity and high dimensionality, our framework uses a special class of statistical distributions that can handle an excess degree of zeros. Although the focus of this study has been based on scRNA-sequencing data generated through droplet-based methods, we show that zero-inflation is required for a significant number of genes to ensure the best distribution fit depending on the nature of the experiment, because around 5% to 20% of the genes under different experimental conditions were detected to follow a zero-inflated distribution. Our study also validates the fact that zero-inflation may be interpreted from a biological perspective i.e. based on cell type membership, which is in line with the findings of Choi *et al*. [18] and Clivio *et al*. [33], and not completely a result that is just an artefact of technical variability.

When compared to other popular tools for identifying differential distributions like *scDD* [12], it is valuable to acknowledge that the *scShapes* approach has the flexibility to account or adjust for multiple confounders in the form of covariates in the GLM. Our approach does not require that prior normalization be done for nuisance factors, but rather enables model-based normalization using the GLM. Furthermore, our method also has the flexibility to perform multiple comparisons across biological conditions and is not restricted to only pair-wise comparisons. This feature provides the option to investigate both global (e.g. effects treatment groups) and pairwise changes (e.g. young versus old), similar to an ANOVA test followed by a post-hoc pairwise comparison but for assessing differential distributions.

One of the tools that exists for identifying the most appropriate statistical model is *M3S* [45], which selects the best fit model from a pool of 11 models using a Kolmogorov-Smirnov test. The 11 distributions considered by *M3S* includes the four distributions included in our *scShapes* method and additional statistical distributions such as Gaussian mixture models. However, we believe that the use of Gaussian distributions to model count data is not advisable as we run the risk of losing information by fitting continuous distributions to count data. Additionally, the use of 11 different distributions makes it more challenging to biologically interpret the best fit distribution for each gene. Hence, for simplicity and efficiency, the four distributions Poisson, NB, ZIP and ZINB represent a sufficient selection for modelling scRNA-seq data.

*scRATE* is another recently published tool for modelling droplet-based scRNA-seq data that uses a mix of unimodal and zero-inflated distributions[18]. This method relies on Bayesian model selection to identify the best model for each gene, and is similar to our method as it also models gene expression distributions using Poisson, NB, ZIP and ZINB distributions using a GLM with an offset term to account for differences in sequencing depth. Importantly, where this method differs from our approach is in its handling of model selection, which is based on the expected log predictive density (ELPD) score. Also the main focus of this study was to identify the origins of zero-inflation rather than to detect differentially distributed genes. Since the key objective of our approach is to detect differential distributions for the purposes of understanding biological perturbations, we ensure that the best fit distribution is identified as appropriate to each gene from the pool of four distributions considered by first performing a Kolmogorov-Smirnov test to identify genes belonging to the family of ZINB distributions. Only the genes that pass through the KS test, would then be considered for the four distributions Poisson, NB, ZIP and ZINB to identify the distribution of best fit. Once model selection is done using the BIC value, we also perform additional model adequacy tests such as testing for the presence of zero-inflation in the genes detected to be following a zero-inflated distribution to ensure that the model distribution is identified correctly and appropriately. The use of the KS test together with the model adequacy tests are crucial since our primary focus is identifying genes that switch distribution shape. It is noteworthy that both the tools *M3S* and *scRATE* fail to first subset genes that belong to the family of statistical distributions under study, but rather assume that all the genes would follow one of the distributions from the pool of distributions considered. Our study, as well as others, present compelling evidence that this assumption is limited or incorrect, especially for scRNA-seq data.

Compared to standard methods of DE, which aim to detect a shift in the mean of gene expression data such as *edgeR*, our method does not make the assumption that all the genes in the transcriptome follow a single distribution. While these methods perform well at detecting statistically significant differences in mean between conditions, they fail to detect subtle changes in gene expression levels that do not involve a change in mean. In addition, even though the negative binomial distribution is a flexible distribution that is capable of handling over-dispersion in the data, it lacks the flexibility to explicitly model both structural zeros and random zeros in the data. As shown by Choi *et al*. [18], the zeros we observe in scRNA-seq data are biological in nature and not purely produced by technical artefacts. However, a negative binomial distribution assumes that the zeros are a result of random sampling and does not allow for the recognition of true biological zeros in the data. This assumption that is made by the NB distribution goes against one of the core truths in biology and one of the key strengths of single cell profiling techniques. Hence it is vital that we model gene expression distributions at the gene level to allow for both random zeros and structural zeros in the data to be identified.

Our framework *scShapes* is built on the assumption that, once all sources of variation other than true biological variation are accounted for, the change in distribution we see is due to the phenotypic changes under consideration in the experimental study. Furthermore, our framework does not aim to cluster cells into subtypes as genes are modelled and evaluated independently. We acknowledge that while this framework is computationally expensive and can be time consuming, especially when applied to datasets with large number of cells, it is possible that improvements through parallelization or more advanced usage of high performance computer clustering could address this limitations.

Some of the genes and pathways discovered through our framework overlap with those identified through methods of transcriptomic data where standard assumptions have been made, indicating the framework’s ability to identify known markers. Importantly this method is able to identify additional genes and pathways both at the tissue level and cell specific level, which may point to new knowledge such as in the example of aging. Most importantly the *scShapes* framework has the flexibility to adjust for any number of covariates (e.g. batch effects, cell-types etc.) and perform multiple comparisons across any set biological conditions of interest. Hence modelling a gene’s expression distribution shape at the gene level, provides an opportunity to extract precise genes and pathways not necessarily put forth by traditional methods of DE.

## Methods

### Code Availability

We have implemented our model selection pipeline as an open-source R package, *scShapes*, available from Bioconductor for Linux, Windows and MacOS. The source code is available on github.com/Malindrie/scShapes under a GPL-3 license. Documentation on the functions and examples for *scShapes* are also available on our GitHub repository. Using *scShapes* requires an input of a matrix of UMI counts with genes as rows and cells as UMIs, along with a data frame of covariates to be included in the GLM and a numeric vector of library sizes. The library sizes are calculated as the total UMI counts per cell, which will be used as an offset in the GLM to account for differences in the sequencing depth. We recommend only a few genes be fitted on a local machine due to the computational demand. Hence, we have containerized the software using both docker and singularity to facilitate its use on high-performance computing clusters or with cloud computing. All analysis scripts used in the paper can be accessed at https://github.com/Malindrie/paper-scShapes.

### Methodology

In order to ensure that only expressed genes are included in the analysis, we retained genes expressed (defined as having a read count > 0) in at least 10% of all cells under each treatment condition. We then restricted our attention to the subset of genes common between all three conditions for downstream analysis. The UMI counts for a given gene are modelled using generalized linear models (GLMs) under each condition separately. The sum of all counts in each cell is used as a proxy to normalize the cells for sequencing depth differences. This cell attribute is used as an offset in a regression model along with any other covariates to account for known sources of cell-to-cell variability. The four models being compared are the Poisson (P), Negative Binomial (NB), Zero Inflated Poisson (ZIP) and Zero Inflated Negative Binomial (ZINB), which are special cases of one another, and all belong to the family of ZINB distributions. Hence, only the subset of genes belonging to the ZINB family of distributions is selected via a Kolmogorov–Smirnov (KS) goodness of fit test, with P-values computed through Monte Carlo simulation. For the significant genes from the KS test under each condition, the expression level of a given gene is then modelled using the error distributions Poisson, NB, ZIP and ZINB with a log link function.

### Parameter Estimation

For model fitting and parameter estimation, the R package **pscl** [46] which allows for the fitting of zero-inflated models was used. The two unimodal distributions, Poisson and NB are implemented in R by the *glm* function [47] in the **stats** package and the *glm.nb* function [48] in the **MASS** package respectively. The package **pscl** provides an extension over the existing R packages for fitting GLM, where the regression coefficients are estimated by maximum likelihood estimation (MLE) using the iterative weighted least squares (IWLS) algorithm.

The parameter estimates are obtained by maximizing the likelihood or log likelihood of the parameters for the observed data. The log-likelihood for the observation *y* given the parameters *θ*, expressed as a function of the mean-value parameter, *μ* = *E*(*Y*) is given by;

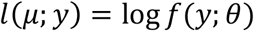

where *f*(*y; θ*) is the density function of *y*. The log-likelihood based on a set of independent observations *y*_1_,…, *y_n_* is the sum of the individual contributions so that,

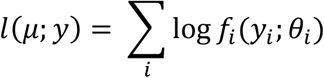

where *μ* = *μ*_1_,…, *μ_n_*

### Model Selection

A critical step in the model building process is the selection of the model that best fits the data among a set of competing models. Among numerous criteria available for model selection, we decided to select the best model based on the least Bayesian Information Criterion (BIC) value as the BIC generally introduces a higher penalty for models with a more complicated parametrization and BIC [49] values are calculated as follows;

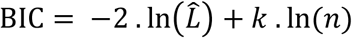

where *k* is the number of free parameters to be estimated, 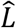 is the maximum value of the likelihood of the model, and *n* is the number of observations.

However, most applications in the literature use different diagnostics in conjunction with information criterion to arrive at the model that best fits the data. Hence further model adequacy tests were done to validate the models selected based on the least BIC value. The deviance goodness of fit test assess the adequacy of the model by comparing the fitted model with the fitted saturated model. The deviance statistic *D*, also known as the likelihood ratio statistic is defined as;

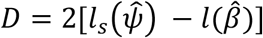

where 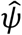 and 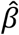 are the MLEs of the saturated and fitted model respectively. The deviance statistics is asymptotically chi-squared distributed with *K* – *p* degrees of freedom, where *K* and *p* are the number of parameters in the saturated and fitted models respectively [50]. i.e.

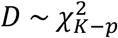

However, the deviance goodness of fit test cannot be used to test the adequacy of the zero-inflated models as these models are not strictly nested i.e. the smaller model sits on the boundary of the parameter space of the larger model [51]. Conversely Molenberghs and Verbeke [17] showed that the resulting likelihood ratio (LR) test statistic of the form;

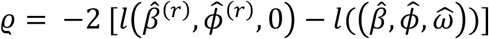

where *ω* is the zero inflation parameter and 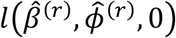 the LR of the model with all parameters of the model estimated with the restriction *ω* = 0, follows an equal mixture of a chi-squared distribution with 0 degrees of freedom and a chi-squared distribution with 1 degree of freedom [51]. i.e.

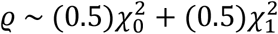

Hence we test for the presence of zero-inflation in the zero-inflated models selected based on the least BIC value using the above test.

### Identifying Differentially Distributed Genes

Each gene is modelled using a GLM where the read count *y_ij_* for gene *i* in cell *j* is assumed to follow one of the following distributions, namely, Poisson, NB, ZIP or ZINB under each biological condition. After performing model adequacy tests detailed above, the distribution that best fits for each gene is identified. Once the statistical distribution for each gene is identified, comparisons in distribution shapes between biological conditions are carried out, to identify genes switching distribution shape between treatment conditions.

### Datasets used

#### Metformin mouse dataset

The scRNA-seq metformin dataset was collected from mice on the two tissues, adipose and muscle. In the study design, three groups of mice, each group consisting of four male C57BL/6J mice, with the young group composed of 3-4 months old mice, old group composed of 18-19 months old mice and treated group composed of 18-19 months old mice treated with 1000ppm (0.1% w/w) metformin for 6 weeks [52]. Using single-cell RNA-Seq, the transcriptome of ~13000 cells from adipose stromal-vascular fraction and ~5000 cells from skeletal muscle were sequenced. The dataset was sequenced on 10X Chromium scRNA-seq platform. We used the pre-processed UMI counts, obtained using Cellranger version 1.3 (10X Genomics). Downstream analysis using Seurat version 3.0.0 [53] identified 12 known cell types in adipose and 13 known cell types in muscle.

#### Simulated T cells dataset

Human T-cells of >50,000 resting and activated cells have been isolated from mucosal and lymphoid sites of two adult deceased organ donors and blood from two healthy donors [43]. The scRNA-seq data were downloaded from the Gene Expression omnibus (GEO) under accession number GSE126030 [https://www.ncbi.nlm.nih.gov/geo/query/acc.cgi?acc=GSE126030]. Similar to the metformin dataset, we applied filters to retain only genes expressed in at least across 10% of the cells to ensure we only included genuinely expressed genes.

#### Peripheral blood mononuclear cells (PBMCs) from patients hospitalized for COVID-19

This dataset consists of scRNA-sequencing data of peripheral blood mononuclear cells (PBMCs) from seven patients hospitalized for COVID-19 (male patients) and six healthy controls (4 male and 2 female) [40]. We used the processed count matrices downloaded from the COVID-19 Cell Atlas (https://www.covid19cellatlas.org/#wilk20) along with the de-identified metadata and embedding. We used information on the 20 cell-types annotated along with the patient ID and gender as covariates in our modelling framework.

#### Simulation study for testing the differential distribution pipeline

To get a realistic set of model parameters, we first performed model classification using our pipeline on the 3k-cell PBMC dataset to learn parameters of the four distributions for each gene. The 3k PBMCs were blood samples from one healthy donor generated using the v1. chemistry and preprocessed with CellRanger 1.1.0. [54] (https://support.10xgenomics.com/single-cell-gene-expression/datasets/1.1.0/pbmc3k). We then generated UMI counts using parameters estimated for each gene by sampling a total cell UMI count from the input data. Using the model classification and parameter estimation we simulated count data for three sample sizes, 2638, 3k and 5k cells. The scRNA-seq counts were generated using the R functions *rpois, rnegbin, rzipois, rzinegbin* available through the R packages **stats**, **MASS** and **VGAM** respectively. We then applied our framework on the simulated data and checked for summary statistics such as accuracy, sensitivity and specificity.

Accuracy of the simulation study reflects the ability of *scShapes* to differentiate between the four different distributions considered. This is calculated as:

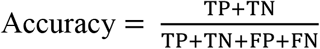

Sensitivity of the simulation study reflects the ability of *scShapes* to determine the correct distribution or the true positives. This is calculated as:

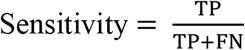

Specificity of the simulation study reflects the ability of *scShapes* to determine the true negatives. This is calculated as:

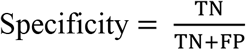

Where for a given gene which is known to be following a Poisson distribution

True positive (TP) = the number of cases correctly identified as Poisson

False positive (FP) = the number of cases incorrectly identified as Poisson

True negative (TN) = the number of cases correctly identified as NB, ZIP, ZINB

False negative (FN) = the number of cases incorrectly identified as NB, ZIP, ZINB

#### Evaluating the significance of overlapping gene sets

To test the significance of overlapping sets of genes, when applying *scShapes* with and without accounting for known sources of biological variability, we used the R package *SuperExactTest* [55], both for calculating the significance of overlap between the gene sets and for visualizations.

#### Pathway over-representation analysis

Pathway over-representation analysis was done using the MSigDB – GSEA online tool by the Broad Institute (https://www.gsea-msigdb.org/gsea/msigdb). This tool calculates the overlap between user provided gene sets and that of the MSigDB collection of datasets chosen by the user. An estimate of the statistical significance of the overlapping gene set is provided based on the hypergeometric P-value which is corrected for multiple hypothesis testing using Benjamini and Hochberg method. All overlaps significant at *p-value* < 0.05 were selected with visualization of top significant pathways. Visualization of the significant pathways was done using the package *ggplot2* in R version 4.0.2.

#### Methods comparison between scShapes, edgeR and scDD

*edgeR* [37] uses a negative binomial generalized linear model for differential expression testing. The quasi-likelihood pipeline using the function *glmQLFTest* implemented in *edgeR* was used for identifying the differentially expressed genes in the pair-wise comparisons of interest, as this method allows for stricter error control than other pipelines of *edgeR*. Lowly expressed genes were filtered (expressed in less than 10% of the cells) and the mouse ID and cellular detection rate (average number of genes expressed per cell) used as covariates to account for the differences in the mice and sequencing depth respectively. Genes were corrected using the Benjamini & Hochberg (BH) correction and only genes that had a corrected *P-value* of < 0.01 were considered to be significantly differentially expressed.

*scDD* [12] utilizes a Bayesian modelling scheme to determine genes with differential distributions classifying them into one of four categories, differential unimodal (DU), differential modality (DM), differential proportion (DP); DB (both DM and DU). Similarly lowly expressed genes were filtered and scran Normalization implemented using the *preprocess* function from *scDD*. Based on the output from the function *scDD* genes falling into one of the four categories were considered to be differentially distributed.

#### Assessing the prevalence of housekeeping genes

To overlap genes with housekeeping genes, we used a previously published list of ubiquitous genes from He *et al*. (2020) [19] downloaded from https://github.com/brianpenghe/Matlab-genomics/blob/master/He_2020_ENCODE3_RNA/GeneLists/Bulk%20Cluster%20Ubiquitous.txt.

## Abbreviations

ANOVA: Analysis of variance
BH: Benjamini & Hochberg
BIC: Bayesian Information Criterion
BL: Blood
BM: Bone marrow
DB: Both differential modality and different component means
DC: Dendritic cells
DD: Differential distribution
DE: Differential expression
DM: Differential modality
DP: Differential proportion
DZ: Differential zeroes
ELPD: Expected log predictive density
GLM: Generalized linear model
KS: Kolmogorov-Smirnov
LG: Lungs
LN: Lymph node
LRT: Likelihood ratio test
NB: Negative binomial
NK: Natural killer
PBMC: Peripheral blood mononuclear cells
QLF: Quasi-likelihood approach
scRNA-seq: Single-cell RNA sequencing
ZINB: Zero inflated negative binomial
ZIP: Zero inflated Poisson

## Acknowledgements

The authors like to thank Dr Alan Huang at the School of Mathematics and Physics, The University of Queensland for the constructive feedback given on the *scShapes* framework.

## Funding

This work is supported by an Australian Research Council Future Fellowship (FT170100047) and a Georgina Sweet Award to J.C.M; Australasian Genomic Technologies Association (AGTA) PhD Top-Up Scholarship to M.D.

## Availability of data and materials

The metformin aging data has been deposited in GEO with accession number GSE194386.

Details on additional results are provided in supplement.

## Authors’ contributions

J.C.M., A.T.F. and M.D. formulated the problem. M.D. developed the *scShapes* method and software with input from J.C.M. and A.T.F. M.D designed and implemented the simulations and applied *scShapes* to the case studies on real scRNA-seq data. J.C.M., A.T.F. and M.D. interpreted the results with input from A.S.K. J.C.M., A.T.F. and M.D. wrote the manuscript with input from A.S.K. All authors read and approved the final version of the manuscript.

## Competing interests

The authors declare that they have no competing interests.

## Ethics approval and consent to participate

Not applicable.

## Notes

### Competing Interest Statement

The authors have declared no competing interest.

## References

1. Buettner, F., et al., Computational analysis of cell-to-cell heterogeneity in single-cell RNA-sequencing data reveals hidden subpopulations of cells. Nature Biotechnology, 2015. 33(2): p. 155–160.

2. Mar, J.C., The rise of the distributions: why non-normality is important for understanding the transcriptome and beyond. Biophys Rev, 2019. 11(1): p. 89–94.

3. Nguyen, A., et al., Single Cell RNA Sequencing of Rare Immune Cell Populations. Frontiers in Immunology, 2018. 9(1553).

4. Jackson, C.A., et al., Gene regulatory network reconstruction using single-cell RNA sequencing of barcoded genotypes in diverse environments. Elife, 2020. 9.

5. Cuomo, A.S.E., et al., Single-cell RNA-sequencing of differentiating iPS cells reveals dynamic genetic effects on gene expression. Nature Communications, 2020. 11(1): p. 810.

6. Kharchenko, P.V., L. Silberstein, and D.T. Scadden, Bayesian approach to single-cell differential expression analysis. Nature Methods, 2014. 11(7): p. 740–742.

7. Shalek, A.K., et al., Single-cell transcriptomics reveals bimodality in expression and splicing in immune cells. Nature, 2013. 498(7453): p. 236–240.

8. Torrenté, L.d., et al., The shape of gene expression distributions matter: how incorporating distribution shape improves the interpretation of cancer transcriptomic data. 2019, bioRxiv.

9. Finak, G., et al., MAST: a flexible statistical framework for assessing transcriptional changes and characterizing heterogeneity in single-cell RNA sequencing data. Genome Biology, 2015. 16(1): p. 278.

10. Chen, Y., et al., edgeR: differential expression analysis of digital gene expression data. 2019.

11. Love, M.I., W. Huber, and S. Anders, Moderated estimation of fold change and dispersion for RNA-seq data with DESeq2. Genome Biology, 2014. 15(12): p. 550.

12. Korthauer, K.D., et al., A statistical approach for identifying differential distributions in single-cell RNA-seq experiments. Genome Biology, 2016. 17(1): p. 222.

13. L. Lun, A.T., K. Bach, and J.C. Marioni, Pooling across cells to normalize single-cell RNA sequencing data with many zero counts. Genome Biology, 2016. 17(1): p. 75.

14. Wagner, A., A. Regev, and N. Yosef, Revealing the vectors of cellular identity with single-cell genomics. Nature Biotechnology, 2016. 34(11): p. 1145–1160.

15. Tanay, A. and A. Regev, Scaling single-cell genomics from phenomenology to mechanism. Nature, 2017. 541(7637): p. 331–338.

16. Larsson, A.J.M., et al., Genomic encoding of transcriptional burst kinetics. Nature, 2019. 565(7738): p. 251–254.

17. Molenberghs, G. and G. Verbeke, Likelihood Ratio, Score, and Wald Tests in a Constrained Parameter Space. The American Statistician, 2007. 61(1): p. 22–27.

18. Choi, K., et al., Bayesian model selection reveals biological origins of zero inflation in single-cell transcriptomics. Genome Biology, 2020. 21(1): p. 183.

19. He, P., et al., The changing mouse embryo transcriptome at whole tissue and single-cell resolution. Nature, 2020. 583(7818): p. 760–767.

20. Meng, G. and H. Mei, Transcriptional Dysregulation Study Reveals a Core Network Involving the Progression of Alzheimer’s Disease. Frontiers in aging neuroscience, 2019. 11: p. 101–101.

21. Gorgoulis, V., et al., Cellular Senescence: Defining a Path Forward. Cell, 2019. 179(4): p. 813–827.

22. Acoba, M.G., et al., The mitochondrial carrier SFXN1 is critical for complex III integrity and cellular metabolism. Cell Reports, 2021. 34(11): p. 108869.

23. López-Otín, C., et al., The hallmarks of aging. Cell, 2013. 153(6): p. 1194–1217.

24. Kulkarni, A.S., S. Gubbi, and N. Barzilai, Benefits of Metformin in Attenuating the Hallmarks of Aging. Cell Metab, 2020. 32(1): p. 15–30.

25. Lei, Y., et al., Metformin targets multiple signaling pathways in cancer. Chinese Journal of Cancer, 2017. 36(1): p. 17.

26. Wu, N., et al., Metformin induces apoptosis of lung cancer cells through activating JNK/p38 MAPK pathway and GADD153. Neoplasma, 2011. 58(6): p. 482–90.

27. Hartwig, J., et al., Metformin Attenuates ROS via FOXO3 Activation in Immune Cells. Frontiers in immunology, 2021. 12: p. 581799–581799.

28. Martins, R., G.J. Lithgow, and W. Link, Long live FOXO: unraveling the role of FOXO proteins in aging and longevity. Aging Cell, 2016. 15(2): p. 196–207.

29. Ma, X., et al., The nuclear receptor RXRA controls cellular senescence by regulating calcium signaling. Aging Cell, 2018. 17(6): p. e12831.

30. Kulkarni, A.S., et al., Metformin regulates metabolic and nonmetabolic pathways in skeletal muscle and subcutaneous adipose tissues of older adults. Aging Cell, 2018. 17(2): p. e12723.

31. Lahoute, C., et al., Premature aging in skeletal muscle lacking serum response factor. PLoS One, 2008. 3(12): p. e3910.

32. Kim, T.K., et al., Interferon regulatory factor 3 activates p53-dependent cell growth inhibition. Cancer Lett, 2006. 242(2): p. 215–21.

33. Clivio, O., et al., Detecting Zero-Inflated Genes in Single-Cell Transcriptomics Data. bioRxiv, 2019: p. 794875.

34. Nikopoulou, C., S. Parekh, and P. Tessarz, Ageing and sources of transcriptional heterogeneity. Biol Chem, 2019. 400(7): p. 867–878.

35. Rezaei-Lotfi, S., N. Hunter, and R.M. Farahani, β-Catenin: A Metazoan Filter for Biological Noise? Front Genet, 2019. 10: p. 1004.

36. Kumar, R.M. and J.J. Collins, Making a noisy gene: HDACs turn up the static. Mol Cell, 2012. 47(2): p. 151–3.

37. Robinson, M.D., D.J. McCarthy, and G.K. Smyth, edgeR: a Bioconductor package for differential expression analysis of digital gene expression data. Bioinformatics, 2010. 26(1): p. 139–40.

38. Soneson, C. and M.D. Robinson, Bias, robustness and scalability in single-cell differential expression analysis. Nature Methods, 2018. 15(4): p. 255–261.

39. Andrawus, M., et al., The effects of environmental stressors on candidate aging associated genes. Experimental Gerontology, 2020. 137: p. 110952.

40. Wilk, A.J., et al., A single-cell atlas of the peripheral immune response in patients with severe COVID-19. Nature Medicine, 2020. 26(7): p. 1070–1076.

41. Chernyak, B.V., et al., COVID-19 and Oxidative Stress. Biochemistry. Biokhimiia, 2020. 85(12): p. 1543–1553.

42. da Silva, R.P., et al., Circulating Type I Interferon Levels and COVID-19 Severity: A Systematic Review and Meta-Analysis. Frontiers in Immunology, 2021. 12(1717).

43. Szabo, P.A., et al., Single-cell transcriptomics of human T cells reveals tissue and activation signatures in health and disease. Nature Communications, 2019. 10(1): p. 4706.

44. Safran, M., et al., GeneCards Version 3: the human gene integrator. Database (Oxford), 2010. 2010: p. baq020.

45. Zhang, Y., et al., M3S: a comprehensive model selection for multi-modal single-cell RNA sequencing data. BMC Bioinformatics, 2019. 20(24): p. 672.

46. Zeileis, A., C. Kleiber, and S. Jackman, Regression Models for Count Data in R. 2008, 2008. 27(8): p. 25.

47. Chambers, J., T. Hastie, and D. Pregibon. Statistical Models in S. 1990. Heidelberg: Physica-Verlag HD.

48. Venables, W.N. and B.D. Ripley, Modern Applied Statistics with S. 2010: Springer Publishing Company, Incorporated.

49. Schwarz, G., Estimating the Dimension of a Model. The Annals of Statistics, 1978. 6(2): p. 461–464.

50. McCullagh, P. and J.A. Nelder, Generalized Linear Models, Second Edition. Chapman and Hall/CRC Monographs on Statistics and Applied Probability Series. 1989: Chapman and Hall.

51. Wilson, P. and J. Einbeck, A new and intuitive test for zero modification. Statistical Modelling, 2019. 19(4): p. 341–361.

52. Kulkarni, A.S., Metformin modulates aging in a cell-type-specific manner in mouse muscle and adipose. Manuscript in preparation.

53. Butler, A., et al., Integrating single-cell transcriptomic data across different conditions, technologies, and species. Nature Biotechnology, 2018. 36(5): p. 411–420.

54. Zheng, G.X.Y., et al., Massively parallel digital transcriptional profiling of single cells. Nature Communications, 2017. 8(1): p. 14049.

55. Wang, M., Y. Zhao, and B. Zhang, Efficient Test and Visualization of Multi-Set Intersections. Scientific Reports, 2015. 5(1): p. 16923.

