## Supplementary materials for "scShapes: A statistical framework for identifying distribution shapes in single-cell RNA-sequencing data"

**Supplementary Materials and Methods: scShapes: A statistical framework for identifying distribution shapes in single-cell RNA-sequencing data**

**Supplementary Methods**

**Generalized linear models for scRNA-seq count data**

Generalized linear models (GLM) are an extension of classical linear models [1, 2], which have the flexibility of specifying models with response variables following different statistical distributions. The response variable of a GLM is related to the linear model through a link function such as the identity, log, or inverse. Most of the bioinformatics tools developed for the analysis of RNA-seq data use GLMs to model read counts generated by the RNA-seq experiments. For example, packages like edgeR [3], DESeq2 [4], MAST [5], Monocle3 [6] and sctransform [7] use a GLM model for modelling read counts due to its flexibility with adjusting for covariates and ability to perform multiple comparisons across biological conditions. We classify the models into two categories based on the way each model treats the excess zeros.

- **One part models** – these models cannot accommodate excess zeros, and instead models the data using standard distributions. We use the Poisson and Negative Binomial (NB) models which are classical GLMs for count data, where the NB distribution addresses the over-dispersion in count data.
- **Zero-inflated models** – these models are commonly used for modelling data with excess zeros. Zero-inflated models are a type of finite mixture models [8], where it is assumed that the zeros are as a result of two processes, i.e. the first process generating only zero counts known as structural zeros and the second process generating zero counts from a Poisson or NB distribution, known as the random zeros.

GLMs characterize the relationship between the dependent variable $y_{i}$ ($i$ = 1,2, …, $n$) and a set of regressors $x_{i}$ where the conditional distribution of $y_{i}|x_{i}$ is given by the probability density function;

$f\left( y;\lambda,\phi\right)=\exp\left( \frac{y . \lambda-b(\lambda)}{\phi}+c(y, \phi) \right)$

where $\lambda$ is the canonical parameter and $\phi$ the dispersion parameter. The functions $b(.)$ and $c\left( . \right)$ are the known functions determining which family of distributions is used. Furthermore, $E\left( y_{i}|x_{i} \right)= \mu_{i}= b^{'}(\lambda_{i})$ and $V\left( y_{i}|x_{i} \right)= \phi.b^{''}(\lambda_{i})$ are the conditional mean and variance respectively.

The conditional mean $E\left( y_{i}|x_{i} \right)$ is related to the set of regressors $x_{i}$ via a known link function $g(.)$ given by;

$$g\left( \mu_{i} \right)= x_{i}^{T}\beta$$

where $\beta$ represents the vector of regression coefficients.

We assume that scRNA-seq data follows one of the below distributions and our aim is to identify the distribution that best fits each gene amongst these statistical distributions. Below describes the probability mass function (PMF) and properties of the four distributions considered under the above two categories of models.

1. **Poisson model**

The Poisson GLM is commonly used in modelling RNA-seq data. The PMF of the Poisson distribution with rate parameter $\mu$ is given by;

$$f\left( y;\mu\right)= \frac{e^{-\mu}. \mu^{y}}{y!}$$

where, $E\left( y \right)= \mu$ and $V\left( y \right)= \mu$. The log link function, $g\left( \mu\right)=\log(\mu)$ is used such that the mean and the linear predictors are related through a log-linear relationship, where;

$\mu_{i}=\exp(x_{i}^{T}\beta)$

2. **Negative Binomial model**

The NB distribution has the capability of handling over-dispersed count data through a dispersion parameter $\theta$ and also overcomes the limitation of the Poisson distribution where both mean and variance are equal. The PMF of the NB distribution with shape parameter $\theta$ and mean $\mu$ is given by;

$$f\left( y;\mu\right)= \frac{\Gamma(y+\theta)}{\Gamma\left( \theta\right). y!} . \frac{\mu^{y}.\theta^{\theta}}{{(\mu+\theta)}^{y+\theta}}$$

where $\Gamma(.)$ is the gamma function and the variance function is given by $V\left( \mu\right)= \mu+\frac{\mu^{2}}{\theta}$. Similarly the mean and the linear predictors are related through a log-linear relationship.

3. **Zero-inflated Poisson model**

The PMF of the zero-inflated Poisson model (ZIP) is given by;

$$f\left( y;\mu,\pi\right)= \left\{ \begin{aligned} \pi+\left( 1-\pi\right)e^{-\mu} y=0 \\ \left( 1-\pi\right) \frac{e^{-\mu}. \mu^{y}}{y!} y>0 \end{aligned} \right.$$

where $\pi$ is the probability of observing excess zeros and is usually modelled by a binomial GLM. The mean and variance of the above is given by $E\left( y \right)=(1-\pi)\mu$ and $V\left( y \right)= \mu(1-\pi)(1+\mu\pi)$. The corresponding mean and linear predictor relationship using log link function is given by;

$$\mu_{i}=\pi_{i}. 0+\left( 1-\pi_{i} \right) . \exp(x_{i}^{T}\beta)$$

Note that when $\pi_{i}=0$, the zero-inflated Poisson model collapses in to a standard Poisson model.

4. **Zero-inflated Negative Binomial model**

The PMF of the zero-inflated NB (ZINB) model is given by;

$$f\left( y;\mu,\pi,\alpha\right)= \left\{ \begin{aligned} \pi+\left( 1-\pi\right) \frac{1}{{(1 + \alpha\mu)}^{1/\alpha}} y=0 \\ \left( 1-\pi\right) \frac{\Gamma(y + 1/\alpha)}{\Gamma\left( y+1 \right)\Gamma(\frac{1}{\alpha})} \frac{{(\alpha\mu)}^{y}}{{(1+\alpha\mu)}^{y+ \frac{1}{\alpha}}} y>0 \end{aligned} \right.$$

where the same definitions of parameters defined in the ZIP model and the mean applies to the ZINB model as well. However, the variance of the ZINB model is given by $V\left( y \right)= \mu(1-\pi)[1+\mu(\pi+ \alpha)]$, where $\alpha$ is the dispersion parameter. Compared to the ZIP models, the ZINB model has added flexibility to accommodate over-dispersion arising from both excess zeros and heterogeneity [9].

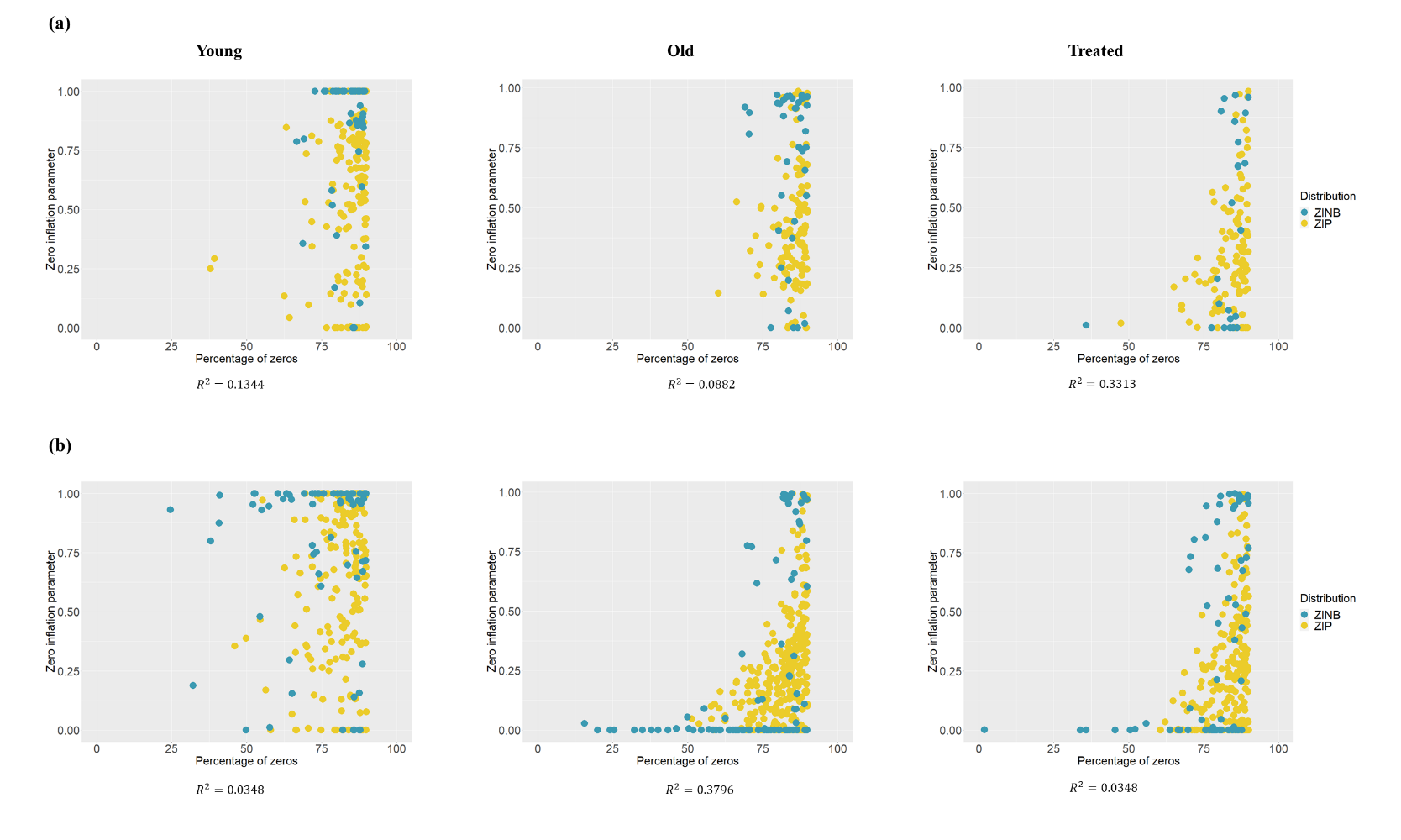
**Supplementary Materials**

**Figure S1: Investigating how the zero inflation parameter is associated with the prevalence of zeroes.** Scatter plot depicts the distribution of zero-inflation parameter against the percentage of zeros in zero-inflated features across all cells in (a) Adipose; (b) Muscle. The zero-inflation parameter π, represents the probability of observing excess zeros in genes that were detected to be following either a ZIP or ZINB distribution and is compared against the percentage of zeros in the corresponding gene.

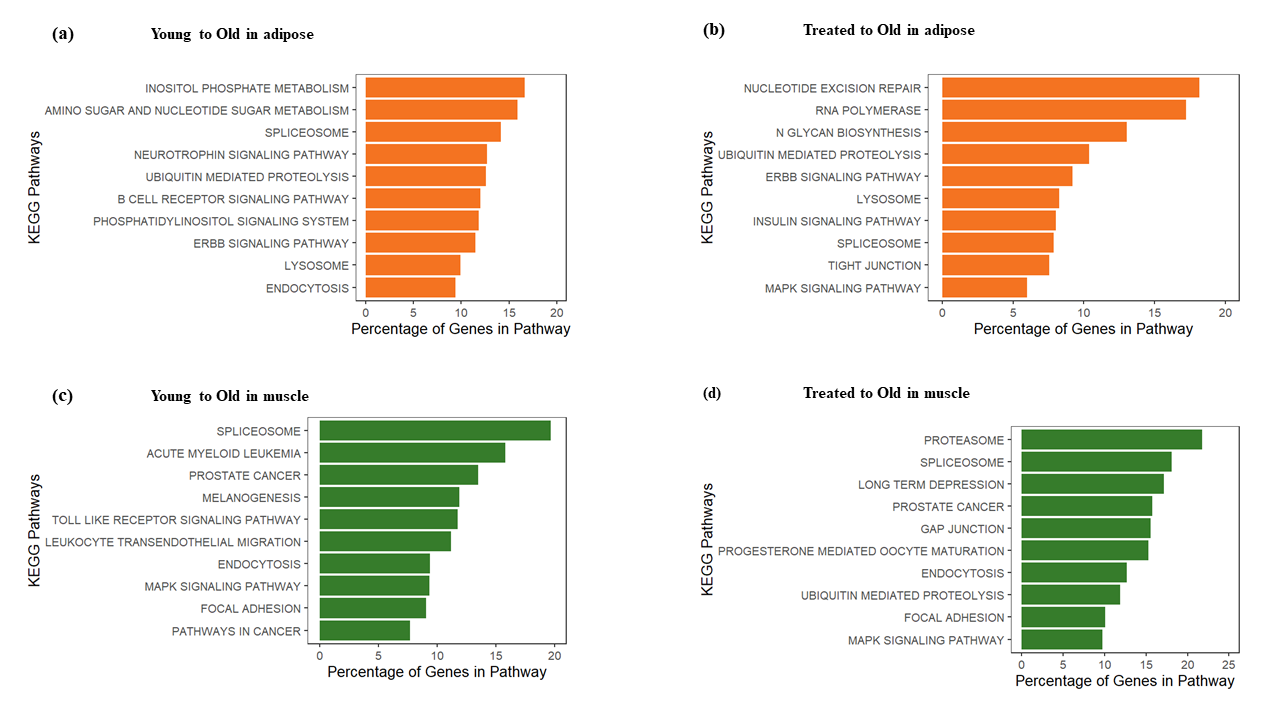

**Figure S2: Pathway over-representation analysis of differentially distributed genes**. The top 10 significant pathways at BH adjusted p-value < $1\times{10}^{-4}$ in the pairwise comparisons (a) Old to Young in adipose (b) Old to Treated in adipose (c) Old to Young in muscle (d) Old to Treated in muscle using KEGG pathways. The length of the bar corresponds to the percentage of differentially distributed genes that overlapped with the curated pathway gene list.

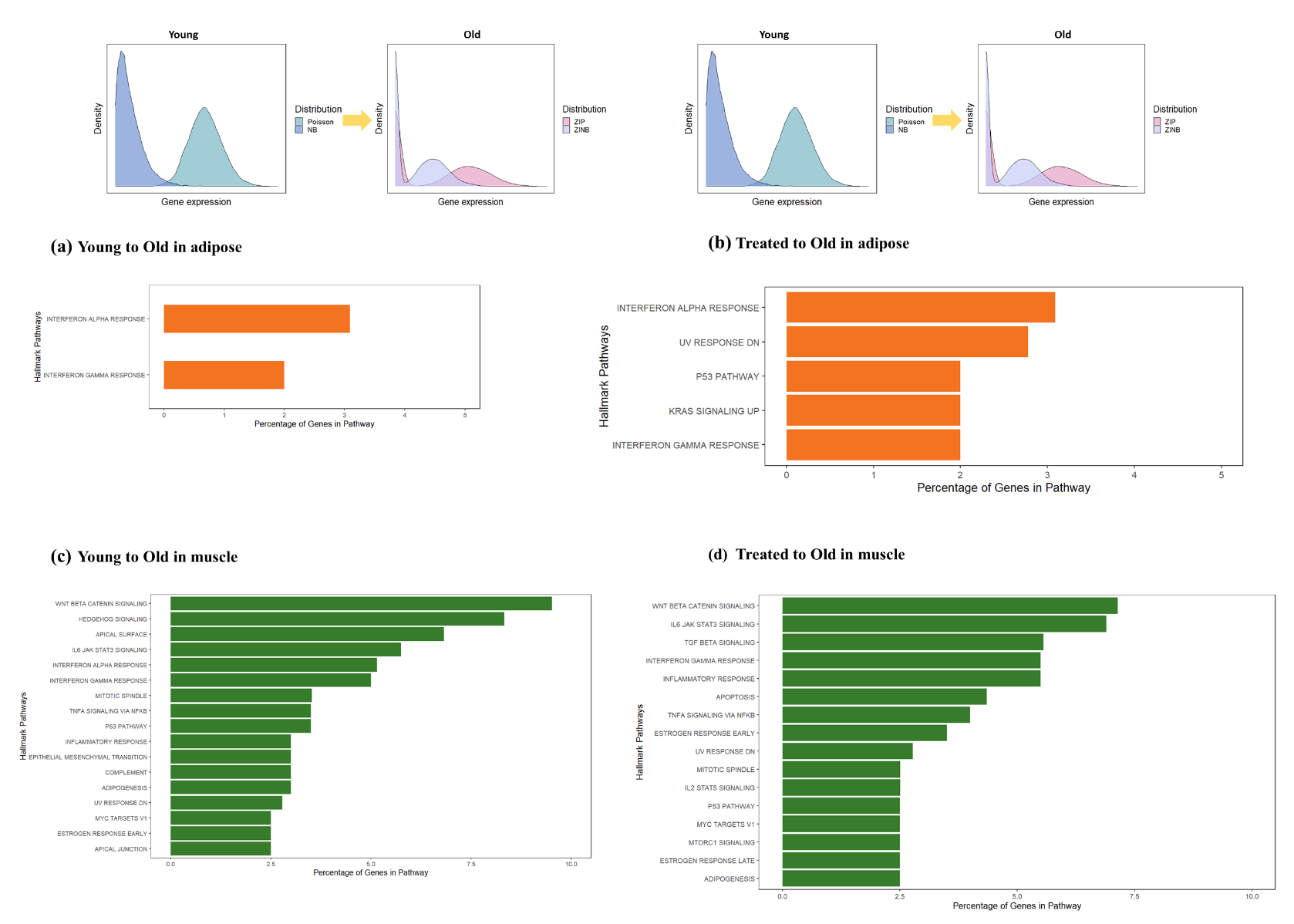
**Figure S3: Pathway over-representation analysis of genes switching between unimodal distributions to zero-inflated distributions**. Genes reverting expression distribution from a Poisson or NB in young/metformin-treated to ZIP or ZINB in old in the pairwise comparisons (a) Young to Old in adipose (b) Treated to Old in adipose (c) Young to Old in muscle (d) Treated to Old in muscle using Hallmark pathways. Significant pathways (BH adjusted p-value < $5\times{10}^{-2}$) are represented where the length of the bar corresponds to the percentage of switching genes that overlapped with the Hallmark gene set.

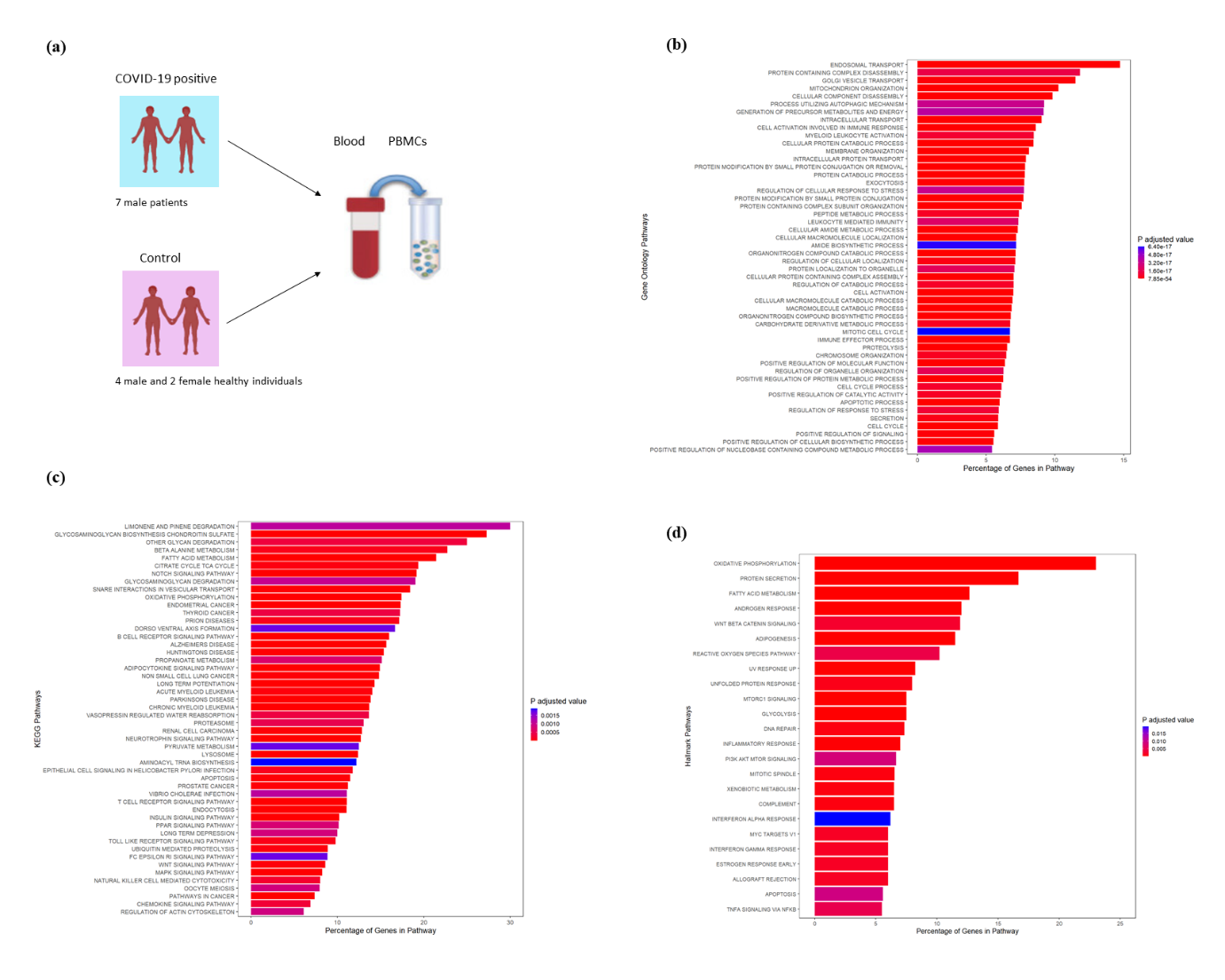
**Figure S4:** (a) Overview of the study design for single-cell RNA-Seq of COVID-19 patients and controls. Single-cells of PBMCs have been profiled from seven male patients hospitalized for COVID19 and four male and 2 female healthy controls. Quality control, normalisation, dimensionality reduction, clustering and differential gene expression analysis has been done using the R package Seurat. By comparing the marker genes per cluster with that of cell-type specific gene from the literature has resulted in 20 cell-types which have been confirmed using the R package *SingleR*. Pathway over-representation analysis of differentially distributed genes between COVID-19 data and healthy controls using (a) GO pathways (b) KEGG pathways and (c) Hallmark pathways. The length of the bar corresponds to the percentage of differentially distributed genes that overlapped with the curated pathway gene list and the colour of the bar corresponds to the BH adjusted P value for the overlap.

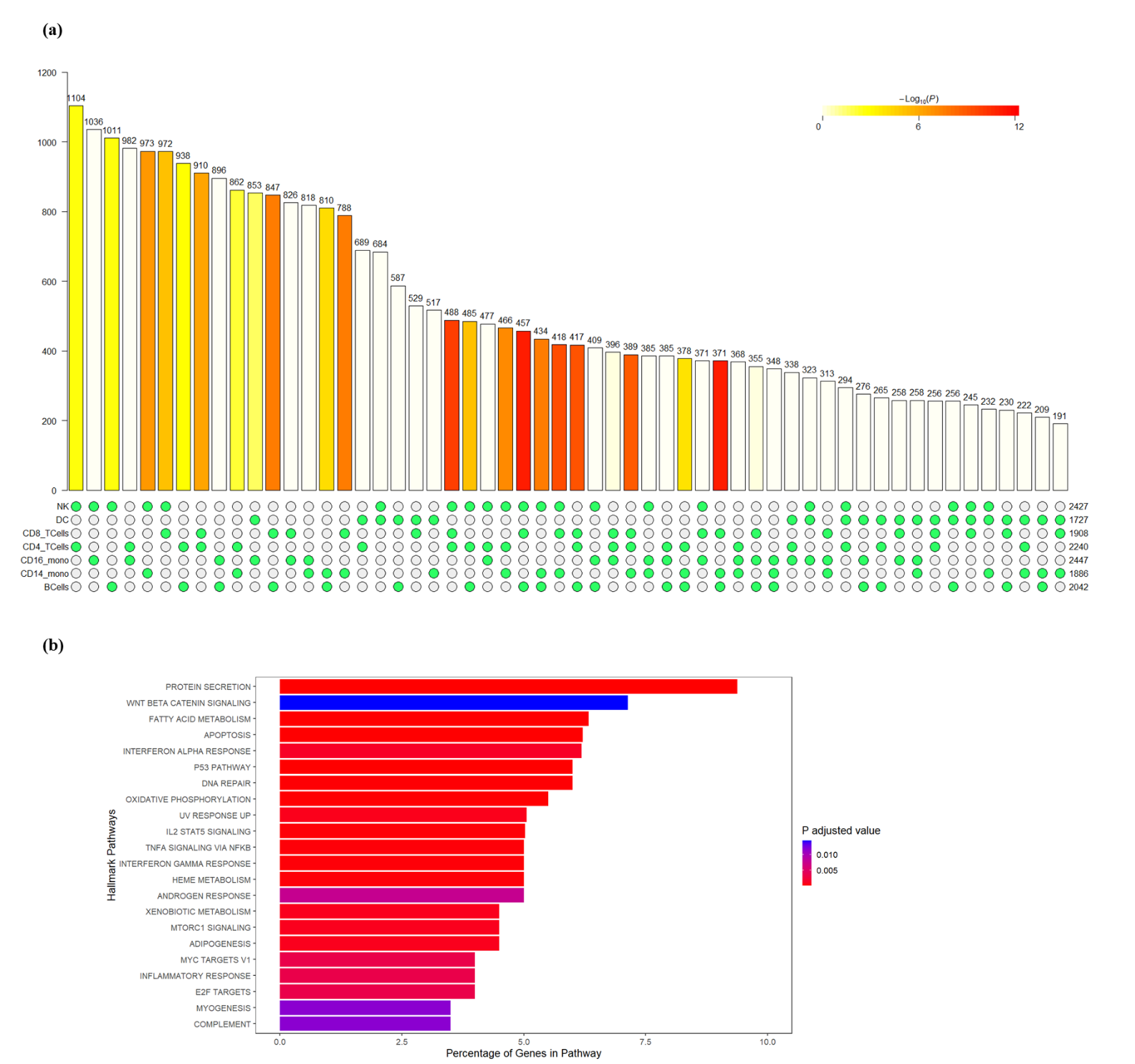
**Figure S5:** (a) Bar plot for overlap of differentially distributed genes at cell-type level between COVID-19 patients and healthy controls. We checked for the COVID-19 driven changes in gene expression distribution at cell type-level and focused our attention on seven main cell-types; Natural Killer cells (NK), B cells, Dendritic cells (DCs), T cells and monocytes. The bar plot summarises the overlap between the genes that switch gene expression distribution between COVID-19 and healthy control groups for different combinations of cell-types under study. The height of each bar corresponds to the number of overlapping genes switching distribution and the colour corresponds to the p-adjusted value for the overlap. (b) Pathway over-representation analysis of commonly differentially distributed genes between NK and T cells. The length of the bar corresponds to the BH adjusted p-value of the overlap of DD genes in NK and T cells. The length of the bar corresponds to the percentage of differentially distributed genes that overlapped with the curated pathway gene list and the colour of the bar corresponds to the BH adjusted P value for the overlap.

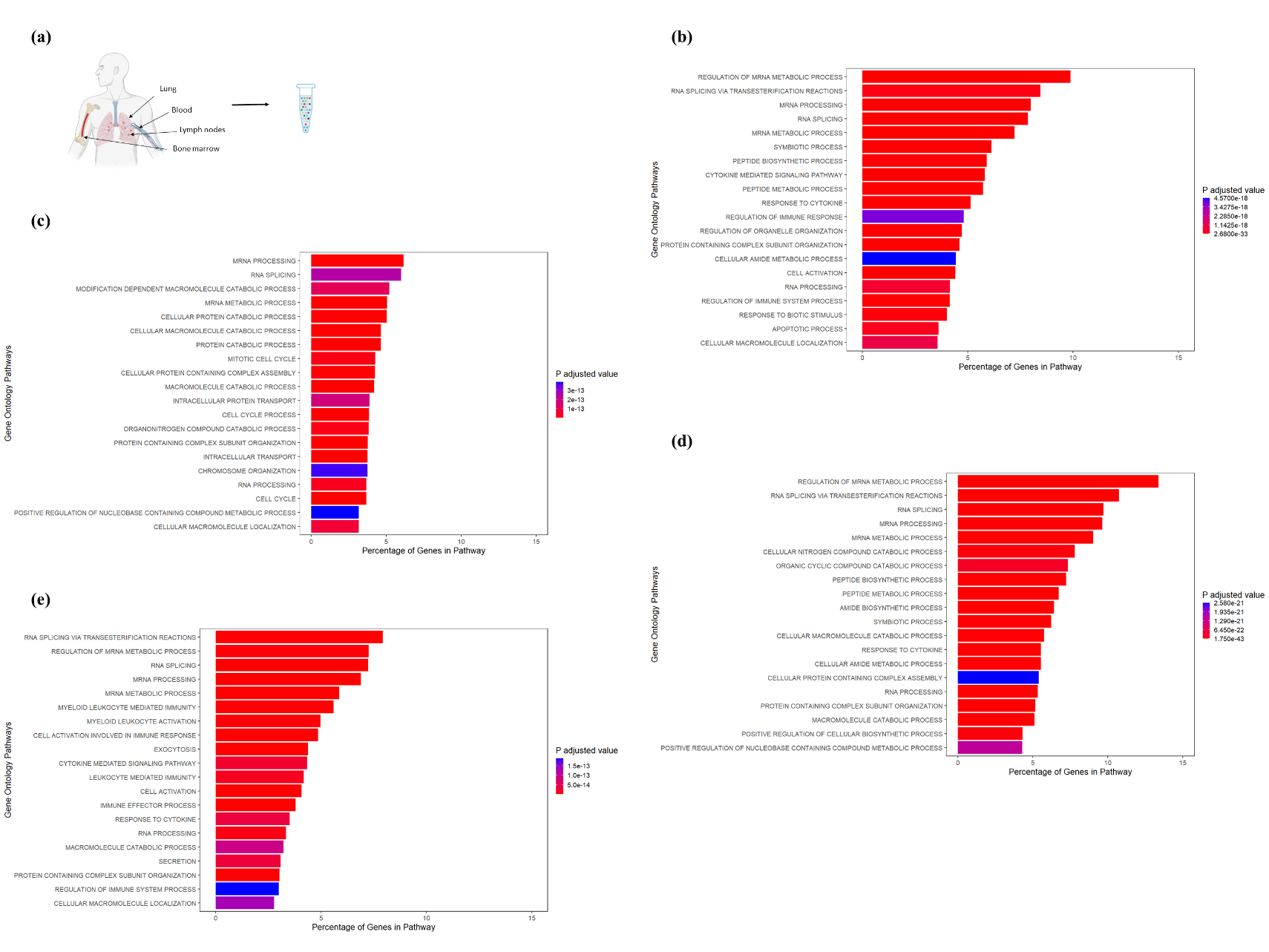
**Figure S6**: (a) Overview of the study design for single-cell RNA-Seq of stimulated T-cells. Single-cells with and without stimulation, have been isolated from lungs, lymph nodes, bone marrow of deceased donors and blood (PBMCs) from healthy donors for comparison. Single-cell sequencing libraries have been sequenced using 10x Chromium Single Cell 3′ Solution. Pathway over-representation analysis of differentially distributed genes in (b) BM (c) BL (d) LG and (e) LN between resting and activated T cells using GO pathways. Here the top 20 significant pathways are visualized. The length of the bar corresponds to the percentage of differentially distributed genes that overlapped with the curated pathway gene list and the colour of the bar corresponds to the BH adjusted P value for the overlap.

**Table S1**: Classification of genes in the Metformin data. Genes were classified as one of four count models (Poisson, NB, ZIP, or ZINB). Table shows the number of genes in each category: (a) using a GLM with the offset term to account for cell sequencing depth and mouse ID as an explanatory covariate; (b) using a GLM that also includes cell type as an explanatory covariate.

(a)

| Tissue | Selected model | Young | Treated | Old |
| --- | --- | --- | --- | --- |
| Adipose | Poisson | 1140 | 1312 | 1353 |
|  | NB | 2830 | 2615 | 2799 |
|  | ZIP | 362 | 547 | 476 |
|  | ZINB | 204 | 462 | 221 |
|  | Total | 4536 | 4936 | 4849 |
| Muscle | Poisson | 1940 | 1703 | 1919 |
|  | NB | 1759 | 1777 | 1841 |
|  | ZIP | 381 | 431 | 324 |
|  | ZINB | 219 | 279 | 137 |
|  | Total | 4299 | 4190 | 4221 |

(b)

| Tissue | Selected model | Young | Treated | Old |
| --- | --- | --- | --- | --- |
| Adipose | Poisson | 1711 | 2051 | 2117 |
|  | NB | 2010 | 1835 | 1840 |
|  | ZIP | 166 | 113 | 124 |
|  | ZINB | 45 | 24 | 40 |
|  | Total | 3932 | 4023 | 4121 |
| Muscle | Poisson | 2305 | 2195 | 2266 |
|  | NB | 1539 | 1444 | 1427 |
|  | ZIP | 208 | 251 | 295 |
|  | ZINB | 66 | 69 | 89 |
|  | Total | 4118 | 3959 | 4077 |

**Table S2**: **Pattern of differential distribution in genes detected as differentially distributed by scShapes.**

(a) Percentage of genes switching distribution from a unimodal to zero-inflated distribution of vice versa

| Tissue | Comparison | Percentage of Switching Genes from Unimodal to ZI (%) |
| --- | --- | --- |
| Adipose | Old vs Young | 27 |
|  | Old vs Treated | 25 |
| Muscle | Old vs Young | 47 |
|  | Old vs Treated | 50 |

(b) Percentage of genes switching distribution from a NB to either Poisson, ZIP or ZINB or vice versa

| Tissue | Comparison | NB to Poisson (%) | NB to ZIP (%) | NB to ZINB (%) |
| --- | --- | --- | --- | --- |
| Adipose | Old vs Young | 72 | 9 | 3 |
|  | Old vs Treated | 75 | 8 | 3 |
| Muscle | Old vs Young | 52 | 13 | 7 |
|  | Old vs Treated | 49 | 12 | 7 |

**Table S3**: Investigating whether switching genes were also detected as differentially-expresssed by edgeR

| Tissue | Comparison | Overlap between DD genes and *edgeR* | Overlap between genes switching from NB to P/ZIP/ZINB and *edgeR* |
| --- | --- | --- | --- |
| Adipose | Old vs Young | 301 | 250 |
|  | Old vs Treated | 125 | 107 |
| Muscle | Old vs Young | 106 | 76 |
|  | Old vs Treated | 89 | 60 |

**Table S4:** Distribution pattern of DZ genes detected by scDD that overlap with the DD genes from scShapes

| Tissue | Comparison | Overlap between DD genes from *scShapes* and DZ from *scDD* | Genes switching from unimodal to ZI distribution or vice versa |
| --- | --- | --- | --- |
| Adipose | Old vs Young | 121 | 36 |
|  | Old vs Treated | 63 | 13 |
| Muscle | Old vs Young | 39 | 18 |
|  | Old vs Treated | 11 | 4 |

**Table S5**: Classification of genes in the COVID-19 data set

| Distribution | Healthy | | COVID-19 | |
| --- | --- | --- | --- | --- |
|  | No of Genes | Percentage | No of Genes | Percentage |
| NB | 3934 | 80% | 3771 | 85% |
| Poisson | 20 | 0% | 5 | 0% |
| ZIP | 423 | 9% | 281 | 6% |
| ZINB | 561 | 11% | 367 | 8% |
| Total | 4938 |  | 4424 |  |

Genes were classified as one of four count models (P, NB, ZIP, or ZINB). Table shows the number of genes in each category using a GLM that includes cell-type, donor ID and gender (in healthy control) as covariates in the model.

**Table S6**: Classification of genes in the human T cells data

| Tissue | Distribution | Rest | | Stim | |
| --- | --- | --- | --- | --- | --- |
|  |  | No of Genes | Percentage | No of Genes | Percentage |
| Lung (LG) | NB | 400 | 24% | 808 | 32% |
|  | Poisson | 1215 | 72% | 1604 | 63% |
|  | ZINB | 12 | 1% | 11 | 0% |
|  | ZIP | 62 | 4% | 124 | 5% |
|  | Total | 1689 |  | 2547 |  |
| Lymph node (LN) | NB | 303 | 15% | 909 | 43% |
|  | Poisson | 1604 | 81% | 1054 | 50% |
|  | ZINB | 6 | 0% | 19 | 1% |
|  | ZIP | 62 | 3% | 118 | 6% |
|  | Total | 1975 |  | 2100 |  |
| Bone marrow (BM) | NB | 245 | 14% | 729 | 32% |
|  | Poisson | 1467 | 82% | 1360 | 60% |
|  | ZINB | 8 | 0% | 16 | 1% |
|  | ZIP | 62 | 3% | 151 | 7% |
|  | Total | 1782 |  | 2256 |  |
| Blood (BL) | NB | 784 | 32% | 1104 | 43% |
|  | Poisson | 1557 | 63% | 1298 | 50% |
|  | ZINB | 15 | 1% | 5 | 0% |
|  | ZIP | 99 | 4% | 178 | 7% |
|  | Total | 2455 |  | 2585 |  |

Genes were classified as one of four count models (P, NB, ZIP, or ZINB). Table shows the number of genes in each category using a GLM that includes donor ID as a covariate in the model.

**Table S7**: Differentially distributed genes between resting and stimulated T cells

| Tissue | No of DD genes |
| --- | --- |
| LG | 368 |
| LN | 515 |
| BM | 438 |
| BL | 456 |

Of the differentially distributed genes between resting and stimulated T cells in the four tissues, a total of 6 genes are DD in all 4 tissues (these include *ETHE1, GSTO1, TSTD1, DUT, USP8, ZBTB7A*).

**References**

1.	McCullagh, P. and J.A. Nelder, *Generalized Linear Models, Second Edition*. Chapman and Hall/CRC Monographs on Statistics and Applied Probability Series. 1989: Chapman and Hall.

2.	Nelder, J.A. and R.W.M. Wedderburn, *Generalized Linear Models.* Journal of the Royal Statistical Society. Series A (General), 1972. **135**(3): p. 370-384.

3.	Robinson, M.D., D.J. McCarthy, and G.K. Smyth, *edgeR: a Bioconductor package for differential expression analysis of digital gene expression data.* Bioinformatics, 2010. **26**(1): p. 139-40.

4.	Love, M.I., W. Huber, and S. Anders, *Moderated estimation of fold change and dispersion for RNA-seq data with DESeq2.* Genome Biology, 2014. **15**(12): p. 550.

5.	Finak, G., et al., *MAST: a flexible statistical framework for assessing transcriptional changes and characterizing heterogeneity in single-cell RNA sequencing data.* Genome Biology, 2015. **16**(1): p. 278.

6.	Qiu, X., et al., *Single-cell mRNA quantification and differential analysis with Census.* Nature Methods, 2017. **14**(3): p. 309-315.

7.	Hafemeister, C. and R. Satija, *Normalization and variance stabilization of single-cell RNA-seq data using regularized negative binomial regression.* Genome Biology, 2019. **20**(1): p. 296.

8.	Cameron, A.C. and P.K. Trivedi, *Regression Analysis of Count Data*. 2 ed. Econometric Society Monographs. 2013, Cambridge: Cambridge University Press.

9.	Rose, C.E., et al., *On the use of zero-inflated and hurdle models for modeling vaccine adverse event count data.* J Biopharm Stat, 2006. **16**(4): p. 463-81.
